# Genus-Wide Genomic Characterization of *Macrococcus*: Insights into Evolution, Population Structure, and Functional Potential

**DOI:** 10.1101/2023.03.06.531279

**Authors:** Laura M. Carroll, Rian Pierneef, Thendo Mafuna, Kudakwashe Magwedere, Itumeleng Matle

**Affiliations:** Department of Clinical Microbiology, SciLifeLab, Umeå University, Umeå, Sweden; Laboratory for Molecular Infection Medicine Sweden (MIMS), Umeå University, Umeå, Sweden; Umeå Centre for Microbial Research, Umeå University, Umeå, Sweden; Integrated Science Lab, Umeå University, Umeå, Sweden; Biotechnology Platform, Agricultural Research Council, Onderstepoort Veterinary Research, Onderstepoort, South Africa; Department of Biochemistry, University of Johannesburg, Auckland Park, South Africa; Directorate of Veterinary Public Health, Department of Agriculture, Land Reform and Rural Development, Pretoria, South Africa; Bacteriology Division, Agricultural Research Council, Onderstepoort Veterinary Research, Onderstepoort, South Africa

**Author notes:** Correspondence: Itumeleng Matle.

**Keywords:** *Macrococcus*, *Macrococcus caseolyticus*, *Macrococcus armenti*, antimicrobial resistance, virulence, cattle, whole-genome sequencing, taxonomy

## Abstract

*Macrococcus* species have been isolated from a range of mammals and mammal-derived food products. While they are largely considered to be animal commensals, *Macrococcus* spp. can be opportunistic pathogens in both veterinary and human clinical settings. This study aimed to provide insight into the evolution, population structure, and functional potential of the *Macrococcus* genus, with an emphasis on antimicrobial resistance (AMR) and virulence potential. All high-quality, publicly available *Macrococcus* genomes (*n* = 104, accessed 27 August 2022), plus six South African genomes sequenced here (two strains from bovine clinical mastitis cases and four strains from beef products), underwent taxonomic assignment (using four different approaches), AMR determinant detection (via AMRFinderPlus), and virulence factor detection (using DIAMOND and the core Virulence Factor Database). Overall, the 110 *Macrococcus* genomes were of animal commensal, veterinary clinical, food-associated (including food spoilage), and environmental origins; five genomes (4.5%) originated from human clinical cases. Notably, none of the taxonomic assignment methods produced identical results, highlighting the potential for *Macrococcus* species misidentifications. The most common predicted antimicrobial classes associated with AMR determinants identified across *Macrococcus* included macrolides, beta-lactams, and aminoglycosides (*n* = 81, 61, and 44 of 110 genomes; 73.6, 55.5, and 40.0%, respectively). Genes showing homology to *Staphylococcus aureus* exoenzyme aureolysin were detected across multiple species (using 90% coverage, *n* = 40 and 77 genomes harboring aureolysin-like genes at 60% and 40% amino acid [AA] identity, respectively). *Staphylococcus aureus* Panton-Valentine leucocidin toxin-associated *lukF-PV* and *lukS-PV* homologs were identified in eight *M. canis* genomes (≥40% AA identity, >85% coverage). Using a method that delineates populations using recent gene flow (PopCOGenT), two species (*M. caseolyticus* and *M. armenti*) were composed of multiple within-species populations. Notably, *M. armenti* was partitioned into two populations, which differed in functional potential (e.g., one harbored beta-lactamase family, type II toxin-antitoxin system, and stress response proteins, while the other possessed a Type VII secretion system; PopCOGenT *P* < 0.05). Overall, this study leverages all publicly available *Macrococcus* genomes in addition to newly sequenced genomes from South Africa to identify genomic elements associated with AMR or virulence potential, which can be queried in future experiments.

## 1 Introduction

Members of the *Macrococcus* genus are Gram-positive, catalase-positive, oxidase-positive, and coagulase-negative cocci (Mazhar et al., 2018; Ramos et al., 2021). The *Macrococcus* genus is a member of the Staphylococcaceae family and was first proposed as a novel genus in 1998, when its four original species (*M. caseolyticus*, *M. equipercicus*, *M. bovicus*, and *M. carouselicus*) were differentiated from members of the closely related *Staphylococcus* genus using numerous genetic and phenotypic characteristics (e.g., 16S rDNA sequencing, DNA-DNA hybridization, pulsed field gel electrophoresis, oxidase activity, cell wall composition, plasmid profiles) (Kloos et al., 1998; Mazhar et al., 2018). Since the four original *Macrococcus* spp. were described in 1998, eight additional *Macrococcus* spp. have been identified (*n* = 12 total validly published *Macrococcus* spp. per the List of Prokaryotic names with Standing in Nomenclature [LPSN], https://lpsn.dsmz.de/genus/macrococcus; accessed 10 December 2022) (Parte et al., 2020): *M. brunensis* (Mannerova et al., 2003)*, M. hajekii* (Mannerova et al., 2003), *M. lamae* (Mannerova et al., 2003), *M. canis* (Gobeli Brawand et al., 2017), *M. bohemicus* (Maslanova et al., 2018), *M. epidermidis* (Maslanova et al., 2018), *M. goetzii* (Maslanova et al., 2018), and *M. armenti* (Keller et al., 2022).

*Macrococcus* spp. have historically been viewed as animal commensals (Mazhar et al., 2018) and have been isolated from a range of mammals (e.g., the skin of cows, pigs, horses, llamas, dogs) and the products derived from them (e.g., dairy products and meat) (Kloos et al., 1998; Mannerova et al., 2003; Cotting et al., 2017; Mazhar et al., 2018; Ramos et al., 2021; Keller et al., 2022). However, the role of *Macrococcus* spp. as opportunistic pathogens has been discussed increasingly in recent years (MacFadyen et al., 2018; Ramos et al., 2021). In veterinary clinical settings, *Macrococcus* spp. have been isolated from infections (e.g., mastitis, otitis, and dermatitis cases, abscesses) in numerous animals, including cattle, sheep, and dogs (Gomez-Sanz et al., 2015; Cotting et al., 2017; Schwendener et al., 2017; Ramos et al., 2021). Notably, in 2018, *Macrococcus* spp. were reportedly isolated from human clinical samples for the first time, when *M. goetzii*, *M. epidermidis*, *M. bohemicus*, and *M. caseolyticus* subsp. *hominis* were isolated from infections at several body sites (i.e., wound sites, gynecological cases, and mycoses cases) (Maslanova et al., 2018). Since then, *M. canis* has additionally been isolated from a human clinical case (i.e., a skin infection) (Jost et al., 2021).

In addition to their pathogenic potential, some *Macrococcus* spp. carry antimicrobial resistance (AMR) genes (Schwendener et al., 2017; MacFadyen et al., 2018; Mazhar et al., 2018; Jost et al., 2021; Ramos et al., 2021). Methicillin resistance in *Macrococcus* spp. is of particular concern, as several mobilizable methicillin resistance determinants (e.g., penicillin-binding protein homologs *mecB*, *mecD*) have been identified in *Macrococcus* spp. (MacFadyen et al., 2018; Mazhar et al., 2018; Ramos et al., 2021). In this context, methicillin-resistant *Macrococcus* strains become particularly concerning: not only can they potentially serve as opportunistic human and veterinary pathogens, but they can potentially transfer mobilizable AMR genes to other organisms, including taxa with a higher virulence potential (e.g., pathogenic *Staphylococcus aureus*) (MacFadyen et al., 2018; Mazhar et al., 2018; Ramos et al., 2021; Schwendener and Perreten, 2022).

Several studies have employed genomic approaches to gain insight into the evolution and population structure of *Macrococcus*; however, these studies relied on a limited number of genomes (Maslanova et al., 2018; Schwendener and Perreten, 2022) and/or focused on specific taxa within the genus (e.g., *M. caseolyticus*) (MacFadyen et al., 2018; Zhang et al., 2022). Furthermore, very few studies—genomic or otherwise—describing *Macrococcus* spp. strains isolated in Africa are available (Tshipamba et al., 2018; Ouoba et al., 2019; Ali et al., 2022). Here, we used whole-genome sequencing (WGS) to characterize six *Macrococcus* spp. strains isolated from bovine-associated sources in South Africa. To gain insight into *Macrococcus* at a genomic scale, we compare our six genomes to all publicly available *Macrococcus* genomes (*n* = 110 total genomes). Overall, our study provides insight into the evolution, population structure, and functional potential of all species—both validly published and putative novel—within the *Macrococcus* genus in its entirety.

## 2 Materials and Methods

### 2.1 Strain isolation

*Macrococcus* strains sequenced in this study were isolated from bovine clinical mastitis specimen sample cases (*n* = 2) and beef products (*n* = 4) and submitted to the Onderstepoort Veterinary Research (OVR) General Bacteriology Laboratory for routine diagnostic services (Supplementary Table S1). From each sample, 10 g (ratio 1:10) were homogenized in buffered peptone water, and then aliquots of 0.1 mL were inoculated onto Baird-Parker agar and Brilliance MRSA 2 agar (both Oxoid, ThermoFisher, Johannesburg) and incubated for 24 hours at 37°C. Presumptive macrococci colonies were streaked onto blood agar supplemented with 5% sheep blood (Oxoid, ThermoFisher, Johannesburg), incubated for 24 hours at 37°C, and identified by phenotypic characteristics as described Poyart et al. (Poyart et al., 2001). Briefly, Gram staining, catalase test, hemolysis, coagulase test, and API 32 ID STAPH (bioMérieux) were used to identify the isolates as macrococci.

### 2.2 Genomic DNA extraction and whole-genome sequencing

Genomic DNA was prepared from overnight cultures using the QIAGEN® DNeasy blood and tissue kit (Germany) according to the manufacturer’s instructions (see section “Strain isolation” above; Supplementary Table S1). WGS of isolates was performed at the Biotechnology Platform, Agricultural Research Council, Onderstepoort, South Africa. DNA libraries were prepared using TruSeq and Nextera DNA library preparation kits (Illumina, San Diego, CA, USA), followed by sequencing on HiSeq and MiSeq instruments (Illumina, San Diego, CA, USA).

### 2.3 Whole-genome sequencing data pre-processing and quality control

Raw Illumina paired-end reads derived from each of the strains isolated here (*n* = 6; see section “Genomic DNA extraction and whole-genome sequencing” above) were supplied as input to Trimmomatic v0.38 (Bolger et al., 2014). Trimmomatic was used to remove Illumina adapters (ILLUMINACLIP:TruSeq3-PE-2.fa:2:30:10:2:keepBothReads), leading and trailing low quality or N bases (i.e., Phred quality < 3; LEADING:3 TRAILING:3), and reads < 36 bp in length (MINLEN:36). FastQC v0.11.9 (https://www.bioinformatics.babraham.ac.uk/projects/fastqc/) was used to evaluate the quality of the resulting trimmed paired-end reads (Supplementary Table S2).

The resulting trimmed paired-end reads associated with each strain were assembled into contigs via Shovill v1.1.0 (https://github.com/tseemann/shovill), using the following parameters (all other parameters were set to their default values): (i) SKESA v2.4.0 (Souvorov et al., 2018) as the assembler (“--assembler skesa”); (ii) a minimum contig length of 200 (“--minlen 200”); (iii) a minimum contig coverage value of 10 (“--mincov 10”). QUAST v5.0.2 (Gurevich et al., 2013) was used to evaluate the quality of each resulting assembled genome (using a minimum contig length parameter of 1 bp), and the “lineage_wf” workflow in CheckM v1.1.3 (Parks et al., 2015) was used to evaluate genome completeness and contamination. MultiQC v1.12 (Ewels et al., 2016) was used to evaluate the quality of all six *Macrococcus* genomes in aggregate (Supplementary Tables S1 and S2).

### 2.4 Acquisition and quality control of publicly available *Macrococcus* spp. genomes

All publicly available GenBank genomes submitted to the National Center for Biotechnology Information (NCBI) Assembly database as members of *Macrococcus* were downloaded (*n* = 102 genomes; accessed 27 August 2022) (Kitts et al., 2016; Schoch et al., 2020). Additionally, all genomes assigned to the *Macrococcus* genus within the Genome Taxonomy Database (GTDB) v207 (Parks et al., 2022), which were not included in the initial set of 102 genomes, were downloaded (*n* = 8 of 88 total GTDB genomes). Together, this search of NCBI and GTDB yielded a preliminary set of 110 publicly available, putative *Macrococcus* genomes.

All 116 putative *Macrococcus* genomes (i.e., 110 publicly available genomes, plus the six genomes sequenced here) were characterized using QUAST and CheckM as described above (see section “Whole-genome sequencing data pre-processing and quality control” above). Six publicly available *Macrococcus* genomes showcased CheckM completeness < 95% and/or QUAST N50 < 20 Kbp; these genomes were excluded from further analysis (*n* = 104 publicly available genomes used in subsequent analyses; Supplementary Table S3). One genome (NCBI GenBank Assembly accession GCA_002119805.1) had >5% CheckM contamination (i.e., 5.11%; Supplementary Table S3). However, because this genome represented the type strain of *M. canis* and was a complete genome, it was used in subsequent steps. Overall, after removing low-quality genomes, the search of NCBI and GTDB, in combination with the six genomes sequenced here, yielded a final set of 110 *Macrococcus* genomes used in subsequent steps (Supplementary Tables S1 and S3).

### 2.5 Taxonomic assignment

All 110 *Macrococcus* genomes (Supplementary Tables S1 and S3; see section “Acquisition and quality control of publicly available *Macrococcus* spp. genomes” above) were assigned to species using the Genome Taxonomy Database Toolkit (GTDB-Tk) v2.1.0 “classify_wf” workflow (default settings) and version R207_v2 of GTDB (Chaumeil et al., 2019; Parks et al., 2022). GTDB-Tk confirmed that all 110 genomes identified here belonged to the *Macrococcus* genus (i.e., either “g Macrococcus” or “g Macrococcus_B”, per GTDB’s nomenclature; these corresponded to the only two GTDB genus designations, which contained the term “*Macrococcus*”; Supplementary Table S4).

Pairwise average nucleotide identity (ANI) values were calculated between all 110 *Macrococcus* genomes using the command-line implementation of OrthoANI v1.40 (Lee et al., 2016) with default settings. The resulting pairwise ANI values were supplied as input to the bactaxR v0.2.1 package (Carroll et al., 2020) in R v4.1.2 (R Core Team, 2021); bactaxR was used to construct a dendrogram and graph of all genomes based on pairwise ANI (dis)similarities, using the ANI.dendrogram and ANI.graph functions, respectively, as well as to construct *de novo* genomospecies clusters using a 95 ANI genomospecies threshold (Supplementary Table S5). OrthoANI was additionally used to calculate ANI values between all 110 *Macrococcus* genomes identified here (query genomes) relative to all *Macrococcus* spp. type strain genomes available in NCBI (reference genomes, *n* = 16 type strain genomes, accessed 4 October 2022; Supplementary Table S3).

Each *Macrococcus* genome was additionally assigned to a marker gene-based species cluster (specI cluster) using classify-genomes (https://github.com/AlessioMilanese/classify-genomes; accessed 3 June 2020) (Milanese et al., 2019) and version 3 of the specI taxonomy (Mende et al., 2013). specI clusters reported by classify-genomes were treated as species assignments (Supplementary Table S6).

The “PopCOGenT” module within PopCOGenT (<u>Pop</u>ulations as <u>C</u>lusters <u>O</u>f <u>Gen</u>e <u>T</u>ransfer, latest version downloaded 31 August 2022) (Arevalo et al., 2019) was additionally used to identify gene flow units among all 110 *Macrococcus* genomes. The resulting “main clusters” reported by PopCOGenT (i.e., gene flow units, which attempt to mimic the classical species definition used for animals and plants) were treated as species assignments (Supplementary Table S7) (Arevalo et al., 2019). Two PopCOGenT main clusters (i.e., Main Clusters 0 and 2; Supplementary Table S7) contained >1 subcluster (i.e., within-species populations identified via PopCOGenT, referred to hereafter as “subclusters”); each of these main clusters was additionally queried individually using the “flexible genome sweeps” module in PopCOGenT to identify subcluster-specific orthologues, using an “alpha” (significance) value of 0.05 (Supplementary Tables S8 and S9) (Arevalo et al., 2019).

### 2.6 *In silico* multi-locus sequence typing

Each of the 110 *Macrococcus* genomes (Supplementary Tables S1 and S3; see section “Acquisition and quality control of publicly available *Macrococcus* spp. genomes” above) was supplied as input to mlst v2.22.0 (https://github.com/tseemann/mlst) for *in silico* multi-locus sequence typing (MLST). Default settings were used so that mlst could auto-select a MLST scheme from PubMLST (Jolley and Maiden, 2010; Jolley et al., 2018). Of the 110 genomes, 62 and 23 genomes were queried using the *M. caseolyticus* (“mcaseolyticus”) and *M. canis* (“mcanis”) PubMLST schemes, respectively; for 25 genomes, no scheme could be applied (Supplementary Table S10).

### 2.7 Genome annotation

Prokka v1.14.6 (Seemann, 2014) was used to annotate each *Macrococcus* genome (*n* = 110, Supplementary Tables S1 and S3; see section “Acquisition and quality control of publicly available *Macrococcus* spp. genomes” above), using the “Bacteria” database and default settings. The “.gff” and “.faa” files produced by Prokka, along with the assembled contigs associated with each strain, were supplied as input to AMRFinderPlus v3.10.40 (Feldgarden et al., 2019), which was used to identify antimicrobial resistance (AMR) determinants in each genome, using the “plus” option (“--plus”, i.e., to enable a search of the extended AMRFinderPlus database, which includes genes involved in virulence, biocide, heat, metal, and acid resistance) and the Prokka annotation format (“-- annotation_format prokka”; Supplementary Table S11).

Amino acid (AA) sequences of virulence factors in the Virulence Factor Database (VFDB) core database (Liu et al., 2019) were downloaded (*n* = 4,188 AA sequences in the VFDB core database; accessed 4 September 2022). CD-HIT v4.8.1 (Li and Godzik, 2006; Fu et al., 2012) was used to cluster all VFDB core database AA sequences using the “cd-hit” command, a sequence identity threshold of 0.4 (“-c 0.4”), and a word length of 2 (“-n 2”, the word size recommended for a 0.4 sequence identity threshold; https://github.com/weizhongli/cdhit/blob/master/doc/cdhit-user-guide.wiki). The “makedb” command in DIAMOND v2.0.15 (Buchfink et al., 2015) was used to construct a DIAMOND database of the VFDB core database in its entirety, and the “diamond blastp” command was used to query AA sequences derived from each *Macrococcus* genome (i.e., “.faa” files produced by Prokka) against the entire VFDB core database, using the following parameters (default values were used for all other parameters): ultra-sensitive mode (“--ultra-sensitive”), one reported maximum target sequence (“-- max-target-seqs 1”, corresponding to the best match produced by DIAMOND: https://github.com/bbuchfink/diamond/issues/29), a minimum percent AA identity threshold of 60% (“--id 60”), and a minimum subject coverage threshold of 50% (“--subject-cover 50”). Each search was repeated using all combinations of (i) minimum percent AA identity thresholds of 0, 40, and 60%, and (ii) minimum subject coverage thresholds of 50 and 90% (Supplementary Tables S12-S17). Because many VFDB virulence factors are composed of multiple genes, and because some genes in VFDB may be highly similar/redundant, virulence factor presence and absence was considered at the whole virulence factor level, where a gene within a given virulence factor was considered to be “present” if any gene within its CD-HIT cluster could be detected in a given genome using DIAMOND. For example, the *Staphylococcus aureus* exotoxin Panton-Valentine leukocidin (PVL) is a two-component toxin (Loffler et al., 2010; Shallcross et al., 2013). In the VFDB core database, PVL (VFDB ID VF0018) is composed of two genes: *lukF-PV* and *lukS-PV* (VFDB IDs VFG001276 and VFG001277, respectively). If any gene within the CD-HIT cluster of *lukF-PV* was detected in a *Macrococcus* genome, *lukF-PV* was considered “present”; likewise, if any gene within the CD-HIT cluster of *lukS-PV* was detected, *lukS-PV* was considered “present”. If both genes were “present”, PVL as a whole was considered to be 100% present. If one gene was “present”, PVL was considered to be 50% present. If neither gene was “present”, PVL was absent (0% present).

Biosynthetic gene clusters (BGCs) were detected in all 110 *Macrococcus* genomes using the command-line implementations of: (i) antiSMASH v6.1.0, using the “bacteria” taxon option (“--taxon bacteria”) and gene finding via Prodigal’s metagenomic mode option (“--genefinding-tool prodigal-m”) (Blin et al., 2021); (ii) GECCO v0.9.2, using the “gecco run” command and the cluster probability threshold lowered to 0.3 (“-m 0.3”; all other settings were set to their defaults) (Carroll et al., 2021). GenBank files (“.gbk”) for all BGCs identified by antiSMASH and GECCO were supplied as input to BiG-SCAPE v1.1.2 (Navarro-Munoz et al., 2020), which was used to cluster the 309 BGCs identified here, as well as experimentally validated BGCs in the MIBiG v2.1 database (“--mibig”) into Gene Cluster Families (GCFs) using default parameter values (Supplementary Table S18) (Kautsar et al., 2020).

### 2.8 Genus-level phylogeny construction

Panaroo v1.2.7 (Tonkin-Hill et al., 2020) was used to identify orthologous gene clusters and construct a core genome alignment (“-a”) among the 110 *Macrococcus* genomes (see section “Acquisition and quality control of publicly available *Macrococcus* spp. genomes” above), plus *Staphylococcus aureus* str. DSM 20231 as an outgroup genome (NCBI RefSeq Assembly accession GCF_001027105.1; *n* = 111 total genomes). The following input/parameters were used (all other parameters were set to their default values): (i) each genome’s “.gff” file produced by Prokka as input (see section “Genome annotation” above); (ii) MAFFT as the aligner (“--aligner mafft”) (Katoh and Standley, 2013); (iii) strict mode (“--clean-mode strict”); (iv) a core genome threshold of 95% (“-- core_threshold 0.95”); (v) a protein family sequence identity threshold of 50% (“-f 0.5”). The core gene alignment produced by Panaroo (“core_gene_alignment.aln”) was supplied as input to IQ-TREE v1.5.4 (Nguyen et al., 2015), which was used to construct a maximum likelihood (ML) phylogeny, using the General Time-Reversible (GTR) nucleotide substitution model (“-m GTR”) (Tavaré, 1986) and 1,000 replicates of the ultrafast bootstrap approximation (“-bb 1000”) (Minh et al., 2013). The resulting ML phylogeny was visualized using the iTOL v6 webserver (https://itol.embl.de/) (Letunic and Bork, 2021).

The genus-level phylogeny produced using Panaroo was compared to genus-level trees constructed using other methods, specifically: (i) PEPPAN (Zhou et al., 2020), a pipeline that can construct pan-genomes from genetically diverse bacterial genomes (e.g., spanning the diversity of an entire genus), and (ii) GTDB-Tk, which, in addition to taxonomic assignment, produces a multiple sequence alignment (MSA) of 120 bacterial marker genes detected in all input genomes (Chaumeil et al., 2019). For (i) PEPPAN, “.gff” files produced by Prokka were used as input (*n* = 111 total genomes, including the *Staphylococcus aureus* outgroup; see section “Genome annotation” above). Default settings were used, except for the “--match_identity” option (the minimal identity of an alignment to be considered during pan-genome construction), which was set to “0.4”, and the “--orthology” option (the algorithm for separating paralogous genes from orthologous genes), which was set to “ml” (i.e., the maximum-likelihood algorithm, reportedly the most accurate) (Zhou et al., 2020). The PEPPAN_parser command was used to produce a Core Genome Allelic Variation (CGAV) tree (using a core genome threshold of 95%; “-a 95”), a gene presence/absence tree (“--tree”), and pan- and core-genome rarefaction curves (“--curve”) (Simonsen et al., 2008; Tettelin et al., 2008; Camacho et al., 2009; Price et al., 2010; Steinegger and Soding, 2017). All aforementioned PEPPAN/PEPPAN_parser steps were repeated three separate times: (a) once as described above, but without the outgroup genome, and (b) using a lower minimal identity threshold (i.e., 20%, “--match_identity 0.2”), with and without the outgroup genome. The resulting trees were annotated using iTOL, and the resulting rarefaction curves were plotted in R using ggplot2 v3.4.0 (Supplementary Figures S1-S6) (Wickham, 2016). For the (ii) GTDB-Tk phylogeny, GTDB-Tk was run as described above, with the addition of the outgroup genome (see section “Taxonomic assignment” above). The resulting AA MSA produced by GTDB-Tk was supplied to IQ-TREE, which was used to construct a ML phylogeny as described above, but with the “LG+F+R4” AA substitution model (i.e., the optimal AA substitution model selected using IQ-TREE’s implementation of ModelFinder, based on Bayesian Information Criteria [BIC] values) (Yang, 1995; Le and Gascuel, 2008; Soubrier et al., 2012; Kalyaanamoorthy et al., 2017). iTOL was used to plot the resulting phylogeny (Supplementary Figure S7).

### 2.9 Functional enrichment analyses

As mentioned above, two PopCOGenT main clusters (i.e., Main Clusters 0 and 2) contained >1 subcluster (see section “Taxonomic assignment” above; Supplementary Table S7). To gain insight into the functional potential of subcluster-specific genes, which had been acquired post-speciation and differentially swept through subclusters identified via PopCOGenT (i.e., flexible genes identified via PopCOGenT, see section “Taxonomic assignment” above; Supplementary Tables S8 and S9), functional enrichment analyses were conducted.

Briefly, for each relevant PopCOGenT main cluster (i.e., Main Cluster 0 and Main Cluster 2; see section “Taxonomic assignment” above), open reading frames (ORFs) produced by PopCOGenT for all members of the given main cluster were supplied as input to the eggNOG-mapper v2.1.9 web server (http://eggnog-mapper.embl.de/; accessed 26 November 2022) (Huerta-Cepas et al., 2019; Cantalapiedra et al., 2021). eggNOG-mapper was used to functionally annotate each ORF, using default settings for all parameters except the input data type option (which was set to “CDS”, as DNA sequences were used as input) and the “Gene Ontology evidence” option, which was set to “Transfer all annotations (including inferred from electronic annotation)”.

For each PopCOGenT subcluster within the given main cluster, enrichment analyses were conducted to identify Gene Ontology (GO) terms (Ashburner et al., 2000; The Gene Ontology Consortium, 2018) assigned via eggNOG-mapper, which were overrepresented among the PopCOGenT flexible genes identified within that particular subcluster: flexible genes identified within the given subcluster were treated as positive instances (PopCOGenT *P* < 0.05; Supplementary Tables S8 and S9), and all other genes within the main cluster were treated as negative instances. Only genes with ≥1 assigned GO term were maintained. GO terms enriched within the positive instances (i.e., the subcluster-specific flexible genes identified via PopCOGenT; Supplementary Tables S8 and S9) were identified via the “runTest” function in the topGO v2.46.0 R package (Alexa et al., 2006), using a Fisher’s exact test (FET) with the “weight01” algorithm. Tests were conducted using each of the Biological Process (BP), Molecular Function (MF), and Cellular Component (CC) ontologies, using a minimum topGO node size of 3 for each ontology (i.e., “nodeSize = 3”, where topGO prunes the GO hierarchy from the terms with < 3 annotated genes). GO terms were considered to be significantly enriched in the flexible genome of a PopCOGenT subcluster if the resulting FET *P*-value was < 0.05; no additional multiple testing correction was applied, as the “weight01” algorithm accounts for GO graph topology and produces *P*-values, which can be viewed as inherently corrected or not affected by multiple testing (Alexa et al., 2006). This approach was repeated for each subcluster within PopCOGenT Main Clusters 0 and 2 (Supplementary Tables S19-S23).

### 2.10 Species-level phylogeny construction

Species-level phylogenies were additionally constructed for the following: (i) GTDB’s *M. caseolyticus* genomospecies, as it was composed of multiple PopCOGenT subclusters and contained five of the six South African genomes sequenced in this study (*n* = 58 genomes; see section “Taxonomic assignment” above); (ii) bactaxR Cluster 13, corresponding to an unknown GTDB genomospecies, which contained the *M. armenti* type strain, because it, too, was composed of multiple PopCOGenT subclusters (*n* = 8 genomes; see section “Taxonomic assignment” above); (iii) bactaxR Cluster 2, as it contained one of the South African genomes sequenced in this study (*n* = 4 genomes; Figure 1).

**Figure 1.**
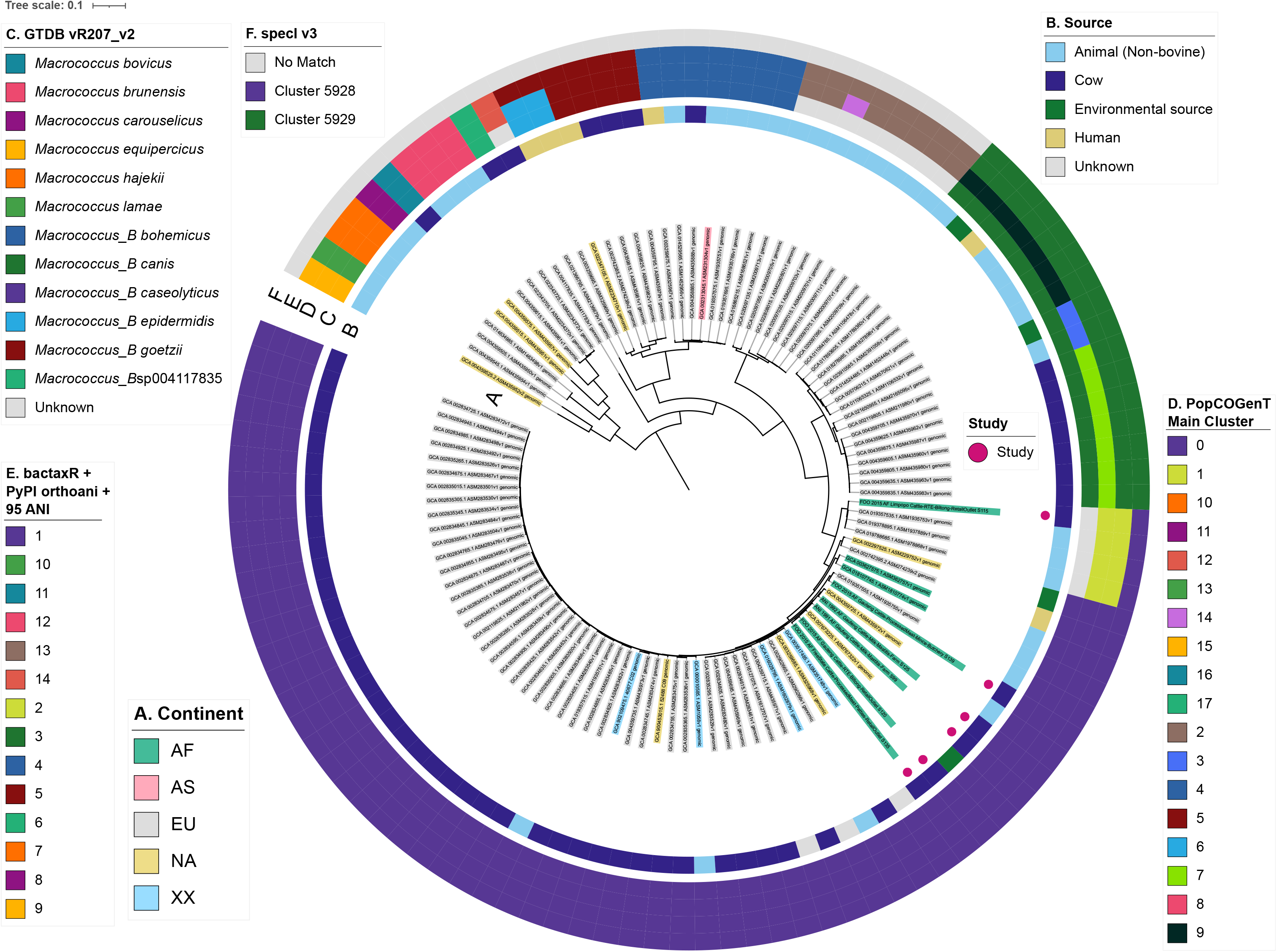
Maximum likelihood (ML) phylogeny of all 104 high-quality, publicly available *Macrococcus* genomes, plus six bovine-associated South African genomes sequenced here (*n* = 110 total *Macrococcus* genomes). Tip label colors (A) denote the continent from which each strain was reportedly isolated. Pink circles denote genomes sequenced in this study (“Study”). Rings surrounding the phylogeny denote (B) the isolation source reported for each strain, as well as (C-F) species assignments obtained using four different taxonomic frameworks: (C) Genome Taxonomy Database (GTDB) species, assigned using the Genome Taxonomy Database Toolkit (GTDB-Tk) v2.1.0 and GTDB vR207_v2; (D) PopCOGenT “main clusters” (i.e., gene flow units, which attempt to mirror the classical species definition for animals and plants); (E) genomospecies clusters delineated *de novo* using average nucleotide identity (ANI) values calculated via OrthoANI, bactaxR, and a 95 ANI genomospecies threshold (i.e., the threshold largely adopted by the microbiological community); (F) marker gene-based species clusters within the specI v3 taxonomy. The ML phylogeny was constructed using an alignment of 649 core genes identified among all 110 *Macrococcus* genomes, plus the genome of *Staphylococcus aureus* str. DSM 20231 (outgroup genome; NCBI RefSeq Assembly accession GCF_001027105.1), using Panaroo and a 50% protein family sequence identity threshold. The tree was rooted using the outgroup (omitted for readability), and branch lengths are reported in substitutions per site. AF, Africa; AS, Asia; EU, Europe; NA, North America; XX, unknown/unreported geographic location.

For *M. caseolyticus* and bactaxR Cluster 13 (i.e., *M. armenti*), Panaroo was used to construct a core gene alignment as described above (see section “Genus-level phylogeny construction” above), using a protein family sequence identity threshold of 70% (“-f 0.7”), all genomes assigned to the respective species cluster as input, and the following outgroup genomes: (i) a *Macrococcus* spp. genome from bactaxR Cluster 2 for *M. caseolyticus* (NCBI GenBank Assembly accession GCA_019357535.1), and (ii) a *M. canis* genome for bactaxR Cluster 13 (NCBI GenBank Assembly accession GCA_014524485.1; Figure 1). Each resulting core gene alignment was supplied as input to IQ-TREE, and ML phylogenies were constructed and annotated as described above (see section “Genus-level phylogeny construction” above).

For *M. caseolyticus*, which was composed of >2 PopCOGenT subclusters, RhierBAPs v1.1.4 (Tonkin-Hill et al., 2018) was additionally employed to cluster the 58 *M. caseolyticus* genomes using two clustering levels. Briefly, Panaroo was used to construct a core gene alignment as described above but with the outgroup genome omitted (*n* = 58 total *M. caseolyticus* genomes). Core SNPs were identified within the resulting core gene alignment using snp-sites v2.5.1 (Page et al., 2016) (using the “-c” option), and the resulting core SNP alignment was supplied as input to RhierBAPs.

For bactaxR Cluster 2, all genomes were fairly closely related (>99.2 ANI via OrthoANI); thus, Snippy v4.6.0 (https://github.com/tseemann/snippy) was used to identify core SNPs among all four genomes within this species cluster, using the closed chromosome of one of the bactaxR Cluster 2 genomes as a reference (NCBI Nucleotide accession NZ_CP079969.1) (Li and Durbin, 2009; Li et al., 2009; Quinlan and Hall, 2010; Li, 2011; Cingolani et al., 2012; Garrison and Marth, 2012; Li, 2013; Tan et al., 2015; Page et al., 2016; Li, 2019; Seemann, 2019). For the bactaxR Cluster 2 genome sequenced in this study, trimmed paired-end reads were used as input; for the three publicly available genomes, assembled genomes were used as input. Snippy was run using default settings, and the resulting cleaned alignment was supplied as input to Gubbins v3.1.3 (Croucher et al., 2015) to remove recombination using default settings. The resulting recombination-free alignment produced by Gubbins was queried using snp-sites as described above, and the resulting core SNP alignment was supplied as input to IQ-TREE. IQ-TREE was used to construct a ML phylogeny using an ascertainment bias correction based on the number of constant sites in the Snippy alignment (“-fconst 645581,381484,377636,655813”), one thousand replicates of the ultrafast bootstrap approximation (“-bb 1000”), and the optimal nucleotide substitution model selected using ModelFinder (“-m MFP”; the K3Pu model, based on its BIC value) (Kimura, 1981; Kalyaanamoorthy et al., 2017). The resulting phylogeny was displayed and annotated using FigTree v1.4.4 (http://tree.bio.ed.ac.uk/software/figtree/). The aforementioned steps were repeated, with the genome sequenced in this study omitted, as the remaining three genomes were highly similar on a genomic scale (>99.99 ANI via OrthoANI for the three publicly available bactaxR Cluster 2 genomes; note that Gubbins was not used here, as there were only three genomes available). Pairwise core SNP distances between genomes were calculated in R using the dist.gene function (with “method” set to “pairwise”) in ape v5.6.2 (Paradis et al., 2004; Paradis and Schliep, 2019). Snippy was additionally used to identify SNPs between other closely related genomes identified in the study (i.e., >99.9 ANI via OrthoANI), using default settings.

## 3 Results

### 3.1 Multiple GTDB species are represented among bovine-associated South African *Macrococcus* strains

Of the *Macrococcus* strains isolated in South Africa that underwent WGS (i.e., two veterinary isolates from bovine clinical mastitis cases, plus four food isolates from beef products), five were assigned to the *M. caseolyticus* genomospecies using the Genome Taxonomy Database Toolkit (GTDB-Tk; Table 1 and Supplementary Table S4). These five genomes each shared 98.0-98.6 average nucleotide identity (ANI) with the closed type strain genome of *M. caseolyticus* (calculated via OrthoANI relative to the *M. caseolyticus* type strain genome with NCBI RefSeq Assembly accession GCF_016028795.1), which is well above the 95 ANI threshold typically used for prokaryotic species delineation (Jain et al., 2018). When compared to each other, the five *M. caseolyticus* genomes sequenced here shared 97.9-99.4 ANI via OrthoANI. One genome (S135) was assigned to PubMLST *M. caseolyticus* sequence type 2 (ST2), while another (S139) was assigned to ST16; the remaining three *M. caseolyticus* genomes belonged to unknown STs (Supplementary Table S10).

**Table 1.** South African Macrococcus spp. genomes sequenced in this study (n = 6).

| Strain | Year of Isolation | Province | Animal | Sample Type | Isolation Source<sup>a</sup> | Establishment Category | GTDB Species<sup>b</sup> |
| --- | --- | --- | --- | --- | --- | --- | --- |
| S99 | 1991 | Gauteng | Cattle | Veterinary clinical sample | Milk from mastitis case | Farm | *M. caseolyticus* |
| S125 | 1992 | Gauteng | Cattle | Veterinary clinical sample | Milk from mastitis case | Farm | *M. caseolyticus* |
| S120 | 2015 | Gauteng | Cattle | Meat sample | RTE beef biltong | Retail outlet | *M. caseolyticus* |
| S139 | 2015 | Gauteng | Cattle | Meat sample | Minced beef | Butchery | *M. caseolyticus* |
| S135 | 2015 | Free State | Cattle | Meat sample | Processed beef patties | Retail outlet | *M. caseolyticus* |
| S115 | 2015 | Limpopo | Cattle | Meat sample | RTE beef biltong | Retail outlet | *M. spp. nov.* |

<sup>a</sup>RTE, ready-to-eat.
<sup>b</sup>Species assigned using the Genome Taxonomy Database (GTDB) Toolkit (GTDB-Tk) v2.1.0 and version R207\_v2 of GTDB; all six genomes were assigned to GTDB’s “*Macrococcus\_B*” genus.

Notably, however, one food isolate (S115) could not be assigned to any known species within GTDB (Table 1 and Supplementary Table S4). Strain S115 was isolated in 2015 from beef biltong, a South African spiced intermediate moisture, ready-to-eat (RTE) meat product, which was being sold in a retail outlet in South Africa’s Limpopo province (Table 1 and Supplementary Table S1). When compared to the five *M. caseolyticus* genomes sequenced here, S115 shared < 95 ANI with each (via OrthoANI). When compared to the type strain genomes of all *Macrococcus* species, S115 was most closely related to *M. caseolyticus* subsp. *hominis* str. CCM 7927 (NCBI RefSeq Assembly accession GCF_002742395.2), sharing 95.3 ANI via OrthoANI. Comparatively, S115 shared 94.6 ANI with the closed *M. caseolyticus* type strain genome (via OrthoANI; NCBI RefSeq Assembly accession GCF_016028795.1).

Overall, these results indicate that, among the bovine-associated South African *M. caseolyticus* genomes sequenced here, (i) considerable within-species diversity exists (e.g., multiple STs are represented, novel STs are present, ANI values between strains sequenced in this study are not particularly high); (ii) one or two *Macrococcus* genomospecies are represented, depending on the species delineation method used (i.e., GTDB or ANI-based comparisons to type strain genomes; Table 1).

### 3.2 Human clinical, veterinary clinical, and food spoilage-associated strains are represented among *Macrococcus* spp. genomes

To compare the bovine-associated South African *Macrococcus* genomes sequenced here to *Macrococcus* genomes collected from other sources in other world regions, the six genomes sequenced here were aggregated with all high-quality, publicly available *Macrococcus* genomes (*n* = 110 total genomes; Figure 1 and Supplementary Table S3). Overall, the complete set of 110 *Macrococcus* genomes represented strains collected from at least ten countries, with most genomes originating from Europe (88 of 110 genomes, 80.0%; Figure 1 and Supplementary Tables S1 and S3).

A vast majority of the genomes (97 of 110 genomes, 88.2%) originated from animal- and animal product-associated sources, with over half of all strains originating from bovine-associated sources (60 of 110 total genomes, 54.5%; Figure 1 and Supplementary Tables S1 and S3). Numerous animal-associated strains, including two strains sequenced here, were reportedly clinical in origin (e.g., isolated from bovine mastitis cases, canine ear infection cases, and wound infections in donkeys; Table 1 and Supplementary Tables S1 and S3). Several animal-associated strains, including four sequenced here, were isolated from food products with the potential for human consumption (i.e., beef and pork meat, cow milk, cheese); one strain was isolated from a food product with a known defect (i.e., “ropy” milk; Table 1 and Supplementary Tables S1 and S3).

Interestingly, six of the 110 *Macrococcus* genomes (5.5%) were derived from human-associated strains (Figure 1 and Supplementary Table S3). At least five of these strains were isolated in conjunction with human clinical cases, including: (i) a hemolytic, methicillin-resistant *M. canis* strain isolated from a 52-year-old immunocompromised patient with cutaneous maculopapular and impetigo lesions (Switzerland, 2019); (ii) a *M. bohemicus* strain from an 80-85 year-old patient with a traumatic knee wound (Czech Republic, 2003); (iii) a *M. goetzii* strain from a foot nail mycosis case in a 30-35 year-old patient (Czech Republic, 2000); (iv) a *M. caseolyticus* subsp. *hominis* strain from an acute vaginitis case in a 40-45 year old patient (Czech Republic, 2003); (v) a *M. epidermidis* strain associated with mycose in a 66-70 year old patient (Czech Republic, 2001; Figure 1 and Supplementary Table S3) (Maslanova et al., 2018; Jost et al., 2021).

Overall, the set of 110 *Macrococcus* genomes aggregated here encompassed strains that were primarily animal- or animal product-associated in origin; however, five strains isolated in conjunction with human clinical cases in Europe were identified (Figure 1).

### 3.3 *Macrococcus* genomospecies clusters may overlap at a conventional 95 ANI threshold

To gain insight into genomic diversity within the *Macrococcus* genus, the following genomospecies delineation methods were applied to the set of 110 *Macrococcus* genomes (i.e., all 104 high-quality, publicly available *Macrococcus* genomes, plus the six genomes sequenced here; Supplementary Tables S1 and S3): (i) GTDB-Tk, a popular genomospecies delineation tool, which relies primarily on a 95 ANI genomospecies threshold; (ii) bactaxR, which uses pairwise ANI values calculated between a set of genomes to delineate genomospecies *de novo* at any user-specified genomospecies threshold (here, ANI values were calculated using OrthoANI, and a 95 ANI genomospecies threshold was used, as this genomospecies threshold has been widely adopted by the microbiological community; Supplementary Figure S8) (Jain et al., 2018); (iii) PopCOGenT (Populations as Clusters Of Gene Transfer) (Arevalo et al., 2019), a method that relies on a metric of recent gene flow to identify species units; (iv) the specI taxonomy, a marker gene-based taxonomic assignment approach (Supplementary Tables S4-S7).

Overall, using GTDB-Tk, *Macrococcus* encompassed 15 genomospecies: 13 defined genomospecies, plus three undefined/putative novel genomospecies defined using a conventional 95 ANI threshold (Figure 1 and Supplementary Table S4). One of these putative novel GTDB-Tk genomospecies encompassed strain S115 sequenced here (denoted as bactaxR Cluster 2 in Figure 1), plus publicly available genomes submitted to NCBI as *M. caseolyticus* (Supplementary Table S3). All members of this genomospecies shared < 95 ANI with the *M. caseolyticus* type strain genome but >95 ANI with the *M. caseolyticus* subsp. *hominis* type strain genome (via OrthoANI, NCBI RefSeq Assembly accessions GCF_016028795.1 and GCF_002742395.2, respectively; Figure 2). The second putative novel GTDB-Tk genomospecies (denoted as bactaxR Cluster 13 in Figure 1) contained the type strain of *M. armenti* (NCBI GenBank Assembly accession GCA_020097135.1); considering *M. armenti* was published as a novel species in 2022, it is likely this genomospecies will be described as such in future versions of GTDB (Keller et al., 2022). The third putative novel GTDB-Tk genomospecies contained a single genome (denoted as bactaxR Cluster 14 in Figure 1), which had been submitted to NCBI as *M. caseolyticus* (NCBI GenBank Assembly accession GCA_021366795.1); however, this genome shared < 75 ANI with the *M. caseolyticus* and *M. caseolyticus* subsp. *hominis* type strain genomes and shared < 81.0 ANI with all other *Macrococcus* spp. genomes (via OrthoANI; Figures 1 and 2).

**Figure 2.**
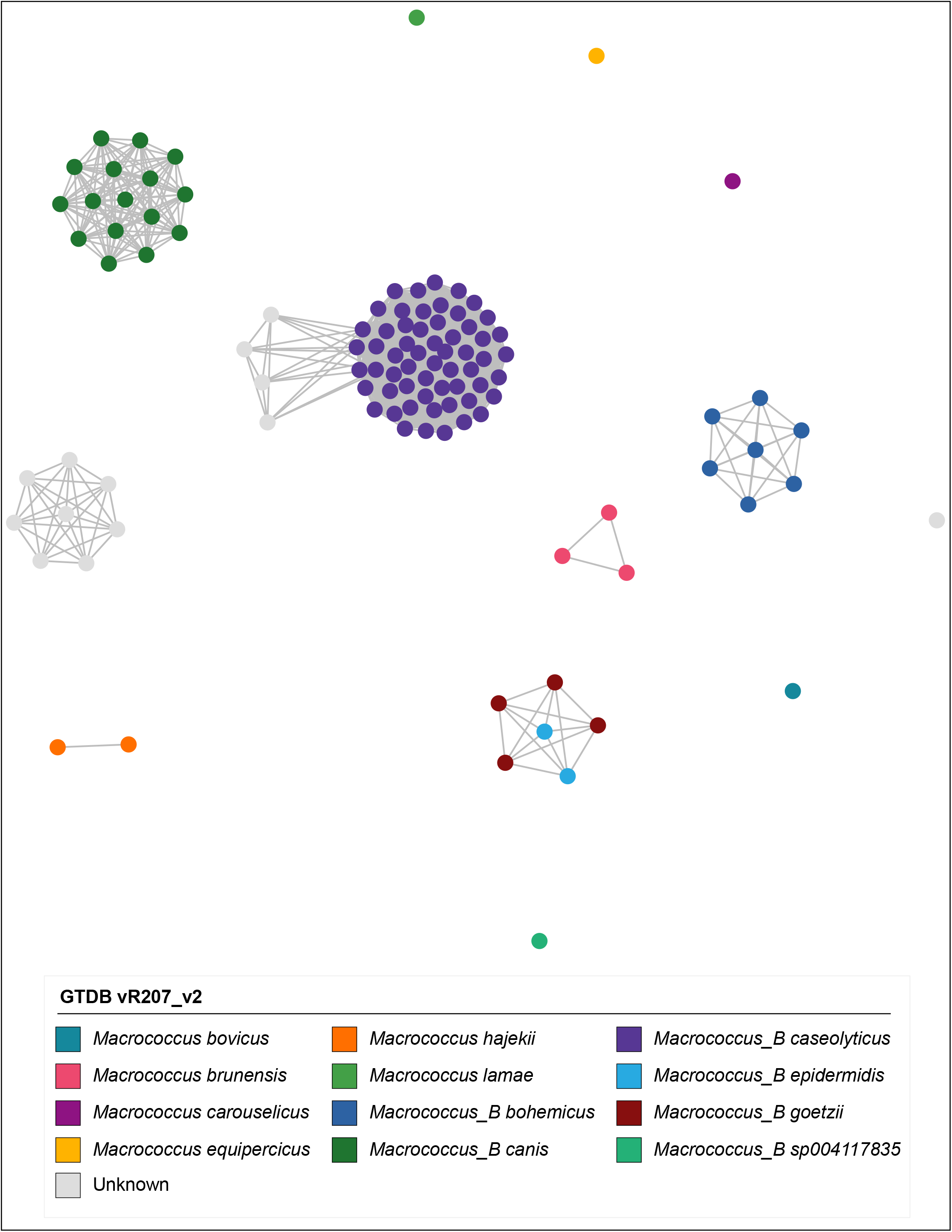
Network constructed using pairwise average nucleotide identity (ANI) values calculated between 110 *Macrococcus* genomes. Nodes represent individual genomes, colored by their Genome Taxonomy Database (GTDB) species assigned via the Genome Taxonomy Database Toolkit (GTDB-Tk) v2.1.0 and GTDB vR207_v2. Two nodes (genomes) are connected if they share ≥95 ANI with each other (calculated via OrthoANI). Networks were constructed and displayed using the ANI.graph function in bactaxR (default settings). See Supplementary Figure S9 for an extended version of this figure, which shows results obtained using all four species delineation methods (i.e., GTDB-Tk, bactaxR with a 95 ANI threshold, PopCOGenT, and specI).

At a conventional 95 ANI threshold, bactaxR produced nearly identical results to GTDB-Tk: 13 of 14 genomospecies defined by bactaxR were identical to those defined by GTDB-Tk, the only difference being that bactaxR aggregated GTDB-Tk’s *M. epidermidis* and *M. goetzii* into a single genomospecies (Figures 1 and 2, Supplementary Figures S8 and S9, and Supplementary Table S5). Similarly, PopCOGenT identified 18 genomospecies among the 110 *Macrococcus* genomes, which were identical to those identified by GTDB-Tk, except: (i) one of the putative novel genomospecies identified by GTDB-Tk and bactaxR was divided into two genomospecies, and (ii) *M. canis* was divided into three genomospecies (Figure 1, Supplementary Figure S9, and Supplementary Table S7). Comparatively, specI identified two defined genomospecies among the *Macrococcus* genomes queried here: (i) Cluster 5928, which encompassed *M. caseolyticus* and a putative novel genomospecies identified by GTDB-Tk, bactaxR, and PopCOGenT, and (ii) Cluster 5929, which was identical to the *M. canis* genomospecies defined by GTDB-Tk and bactaxR (Figure 1, Supplementary Figure S9, and Supplementary Table S6).

Importantly, for three of the four genomospecies delineation methods used here (i.e., GTDB-Tk, bactaxR, and PopCOGenT), genomes assigned to separate genomospecies could share >95 ANI with each other (Figure 2 and Supplementary Figure S9), indicating that some *Macrococcus* genomospecies defined at a conventional 95 ANI threshold overlap. specI did not yield overlapping genomospecies at a conventional 95 ANI threshold (Supplementary Figure S9); however, nearly a third of *Macrococcus* genomes (*n* = 32 of 110 genomes, 29.1%) could not be assigned to a species via specI (Figure 1, Supplementary Figure S9, and Supplementary Table S6).

Taken together, these results indicate that (i) three of the four genomospecies delineation methods queried here (i.e., GTDB-Tk, bactaxR with OrthoANI and a 95 ANI threshold, and PopCOGenT) produced similar, albeit not identical, results when applied to *Macrococcus* (Figure 1); (ii) the same three genomospecies delineation methods produced “overlapping genomospecies”, in which some genomes could share >95 ANI with members of another genomospecies (Figure 2 and Supplementary Figure S9).

### 3.4 Multiple *Macrococcus* spp. contain genomes, which are predicted to be multi-drug resistant

Antimicrobial resistance (AMR) and stress response determinants (detected via AMRFinderPlus; Supplementary Table S11) were variably present throughout *Macrococcus* and were associated with predicted resistance to a variety of antimicrobial classes, heavy metals, and metalloids (Figure 3 and Supplementary Figure S10). The most common classes of antimicrobials for which *Macrococcus* was predicted to harbor resistance determinants included macrolides, beta-lactams, and aminoglycosides (*n* = 81, 61, and 44 of 110 genomes with one or more associated AMR determinants, corresponding to 73.6, 55.5, and 40.0% of *Macrococcus* genomes, respectively; Figure 3, Supplementary Figure S10, and Supplementary Table S11). The high proportion of genomes harboring an ATP-binding cassette subfamily F protein (ABC-F)-encoding gene (*abc-f*) contributed to the high proportion of genomes with predicted macrolide resistance (*n* = 74 of 110 *Macrococcus* genomes harbored *abc-f*, 67.3%), although several additional macrolide resistance genes were sporadically present within the genus (Figure 3, Supplementary Figure S10, and Supplementary Table S11). The high proportion of genomes showcasing predicted beta-lactam resistance, on the other hand, was largely driven by the presence of *mecD* (*n* = 43 of 110 *Macrococcus* genomes, 39.1%), although *mecB* and *bla* were also present in >10% of genomes (Figure 3, Supplementary Figure S10, and Supplementary Table S11). Aminoglycoside resistance genes were sporadically present among *Macrococcus* genomes, the most common being *str* (*n* = 23 of 110 genomes, 20.9%; Figure 3, Supplementary Figure S10, and Supplementary Table S11).

**Figure 3.**
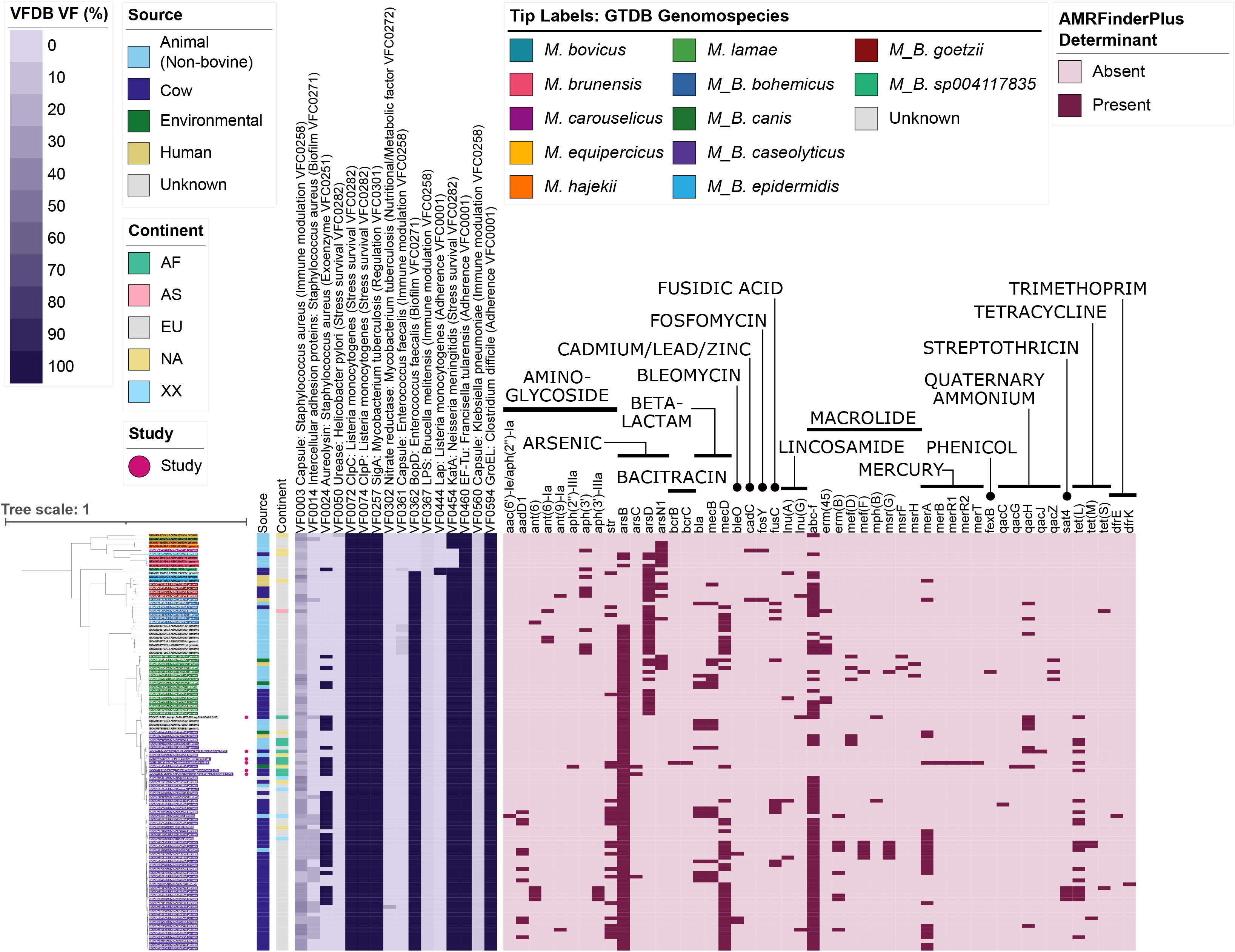
Maximum likelihood (ML) phylogeny of all 104 high-quality, publicly available *Macrococcus* genomes, plus six bovine-associated South African genomes sequenced here (*n* = 110 total *Macrococcus* genomes). Tip label colors correspond to genomospecies assignments obtained via the Genome Taxonomy Database Toolkit (GTDB-Tk) v2.1.0 and GTDB vR207_v2. Genomes of strains isolated and sequenced in this study are denoted by pink circles (“Study”). Color strips and heatmaps to the right of the phylogeny denote (from left to right): (i) the source from which each strain was reportedly isolated (“Source”); (ii) the continent from which each strain was reportedly isolated (“Continent”); (iii) percentage of virulence factors (VF) present in the Virulence Factor Database (VFDB) core database, which were detected in each genome using DIAMOND blastp, with minimum amino acid identity and subject coverage thresholds of 60 and 50%, respectively (“VFDB VF”); (iv) antimicrobial resistance (AMR) and stress response determinants identified in each genome using AMRFinderPlus (default settings; “AMRFinderPlus Determinant”). The ML phylogeny was constructed using an alignment of 649 core genes identified among all 110 *Macrococcus* genomes, plus the genome of *Staphylococcus aureus* str. DSM 20231 (outgroup genome; NCBI RefSeq Assembly accession GCF_001027105.1), using Panaroo and a 50% protein family sequence identity threshold. The tree was rooted using the outgroup (omitted for readability), and branch lengths are reported in substitutions per site. AF, Africa; AS, Asia; EU, Europe; NA, North America; XX, unknown/unreported geographic location.

The most common AMR profiles among *Macrococcus* genomes harboring one or more AMR determinant were those associated with (i) macrolide and (ii) beta-lactam/macrolide resistance (*n* = 18 and 13 of 110 genomes, corresponding to 16.4 and 11.8% of genomes, respectively; Figure 3, Supplementary Figure S10, and Supplementary Table S11). However, numerous predicted multidrug-resistance (MDR) profiles were observed, the most common being (i) aminoglycoside/beta-lactam/macrolide and (ii) aminoglycoside/beta-lactam/macrolide/tetracycline resistance (*n* = 11 and 8 of 110 genomes, corresponding to 10.0 and 7.3% of genomes, respectively; Figure 3, Supplementary Figure S10, and Supplementary Table S11). The genome displaying predicted AMR to the most antimicrobial classes was the genome of *M. caseolyticus* strain 5813_BC74, which had reportedly been isolated from bovine bulk tank milk in the United Kingdom in 2016 (NCBI GenBank Assembly accession GCA_002834615.1; Supplementary Table S3). This genome displayed predicted aminoglycoside/beta-lactam/fusidic acid/lincosamide/macrolide/tetracycline resistance (*n* = 6 antimicrobial classes; Figure 3, Supplementary Figure S10, and Supplementary Table S11).

Predicted AMR phenotypes observed in < 10% of all *Macrococcus* genomes included: (i) fusidic acid resistance (due to the presence of *fusC*; *n* = 10), (ii) lincosamide resistance (per *lnu(A), lnu(G)*; *n* = 7), (iii) streptothricin resistance (via *sat4*; *n* = 4), (iv) bleomycin resistance (via *bleO*; *n* = 3), (v) trimethoprim (via *dfrE, dfrK*) and (vi) fosfomycin resistance (via *fosY*, *n* = 2 genomes each), and (vii) phenicol resistance (via *fexB*, *n* = 1 genome; Figure 3, Supplementary Figure S10, and Supplementary Table S11). Interestingly, one of the South African genomes sequenced here harbored bacitracin resistance genes *bcrB* and *bcrC* (Figure 3, Supplementary Figure S10, and Supplementary Table S11); this strain (i.e., S99, from a bovine mastitis case in Gauteng in 1991) was the only *Macrococcus* genome in which bacitracin resistance genes were detected (Figure 3, Supplementary Figure S10, and Supplementary Table S11).

Overall, these results indicate that (i) numerous AMR determinants are variably present within and among *Macrococcus* species; and (ii) *Macrococcus* genomes may harbor AMR determinants predictive of an MDR phenotype (i.e., resistant to three or more antimicrobial classes; Figure 3, Supplementary Figure S10, and Supplementary Table S11). However, these results should be interpreted with caution, as AMR potential was not evaluated phenotypically.

### 3.5 *Staphylococcus aureus* virulence factor homologues can be detected within some *Macrococcus* genomes at low amino acid identity

To gain insight into the virulence potential of *Macrococcus,* the 110 genomes aggregated here were queried for virulence factors present in the VFDB core database (Figure 3, Supplementary Figure S10, and Supplementary Tables S12-S17). Proteins with homology to stress response- (i.e., *Listeria monocytogenes* ClpC and ClpP, *Neisseria meningitidis* KatA), adherence- (i.e., *Clostridium difficile* GroEL and *Francisella tularensis* EF-Tu), regulatory- (i.e., *Mycobacterium tuberculosis* SigA), and biofilm-associated proteins (i.e., *Enterococcus faecalis* BopD) present in VFDB were detected in over 90% of all *Macrococcus* genomes (i.e., ≥100 of 110 genomes, using minimum amino acid [AA] identity and coverage thresholds of 60% and 50%, respectively; Figure 3, Supplementary Figure S10, and Supplementary Table S16).

Additionally, proteins with homology to immune modulation-associated virulence factor proteins in VFDB (i.e., the *Staphylococcus aureus* and *Klebsiella pneumoniae* capsules, plus the *Brucella melitensis* lipopolysaccharide) were detected in ≥100 of the *Macrococcus* genomes aggregated here (>90% of 110 *Macrococcus* genomes, using minimum AA identity and coverage thresholds of 60% and 50%, respectively; Figure 3, Supplementary Figure S10, and Supplementary Table S16). However, each of these three virulence factors in their entirety could not be detected in any genome, as no more than 40% of the proteins associated with each virulence factor were detected in a single genome (using minimum AA identity and coverage thresholds of 60% and 50%, respectively; Supplementary Table S16).

Several additional proteins showing homology to VFDB virulence factors were variably present within and among *Macrococcus* species (Figure 3, Supplementary Figure S10, and Supplementary Tables S16). Notably, genes encoding the *Staphylococcus aureus* exoenzyme aureolysin could be detected across multiple *Macrococcus* species (using 90% coverage, *n* = 40 and 77 genomes harboring aureolysin-encoding genes at 60% and 40% AA identity, respectively; Figure 3, Supplementary Figure S10, and Supplementary Tables S15 and S17).

Interestingly, using lower AA identity thresholds, proteins showing homology to exotoxin-associated proteins were identified in several *Macrococcus* genomes (Supplementary Figure S10 and Supplementary Tables S12-S17). Most notably, genes sharing homology (i.e., ≥40% AA identity) with *Staphylococcus aureus* Panton-Valentine leucocidin (PVL) toxin-associated *lukF-PV* and *lukS-PV* were identified in eight *M. canis* genomes at >85% coverage (per GTDB-Tk; Supplementary Figure S10 and Supplementary Table S14). For all eight *M. canis* genomes in which they were detected, the *lukF-PV* and *lukS-PV* homologs were located next to each other in the genome (Supplementary Figure S10 and Supplementary Table S14).

Overall, these results indicate that proteins homologous to virulence factors present in other species (e.g., *Staphylococcus aureus*) can be detected in some *Macrococcus* genomes. However, the methods employed here are not adequate to properly evaluate the virulence potential of *Macrococcus* strains that possess these homologs; thus, these results should be interpreted with extreme caution.

### 3.6 *Macrococcus* species differ in pan-genome composition

Using PEPPAN and a 40% AA identity threshold, a total of 10,300 genes were detected among the 110 *Macrococcus* genomes aggregated here, 1,229 of which were core genes present in all 110 genomes (11.9% of all *Macrococcus* genes; Figure 4 and Supplementary Figures S1, S3, and S5). Comparatively, at a 20% AA identity threshold, 9,835 total genes were detected, 1,235 of which were core genes present in all 110 genomes (12.6% of all *Macrococcus* genes; Supplementary Figures S2-S4 and S6). Based on trees constructed using pan-genome element presence/absence, *Macrococcus* species (per GTDB-Tk) tended to cluster together based on pan-genome composition, although not exclusively (Supplementary Figures S1 and S2). Specifically, the topology of the PEPPAN pan-genome tree differed from that of the PEPPAN Core Genome Allelic Variation (CGAV) tree, as some *Macrococcus* species were polyphyletic based on pan-genome element presence/absence (Supplementary Figures S1 and S2). Overall, *Macrococcus* species tend to differ via both core genome phylogeny (Figures 1 and 3) and pan-genome composition (Figure 4 and Supplementary Figures S1-S6).

**Figure 4.**
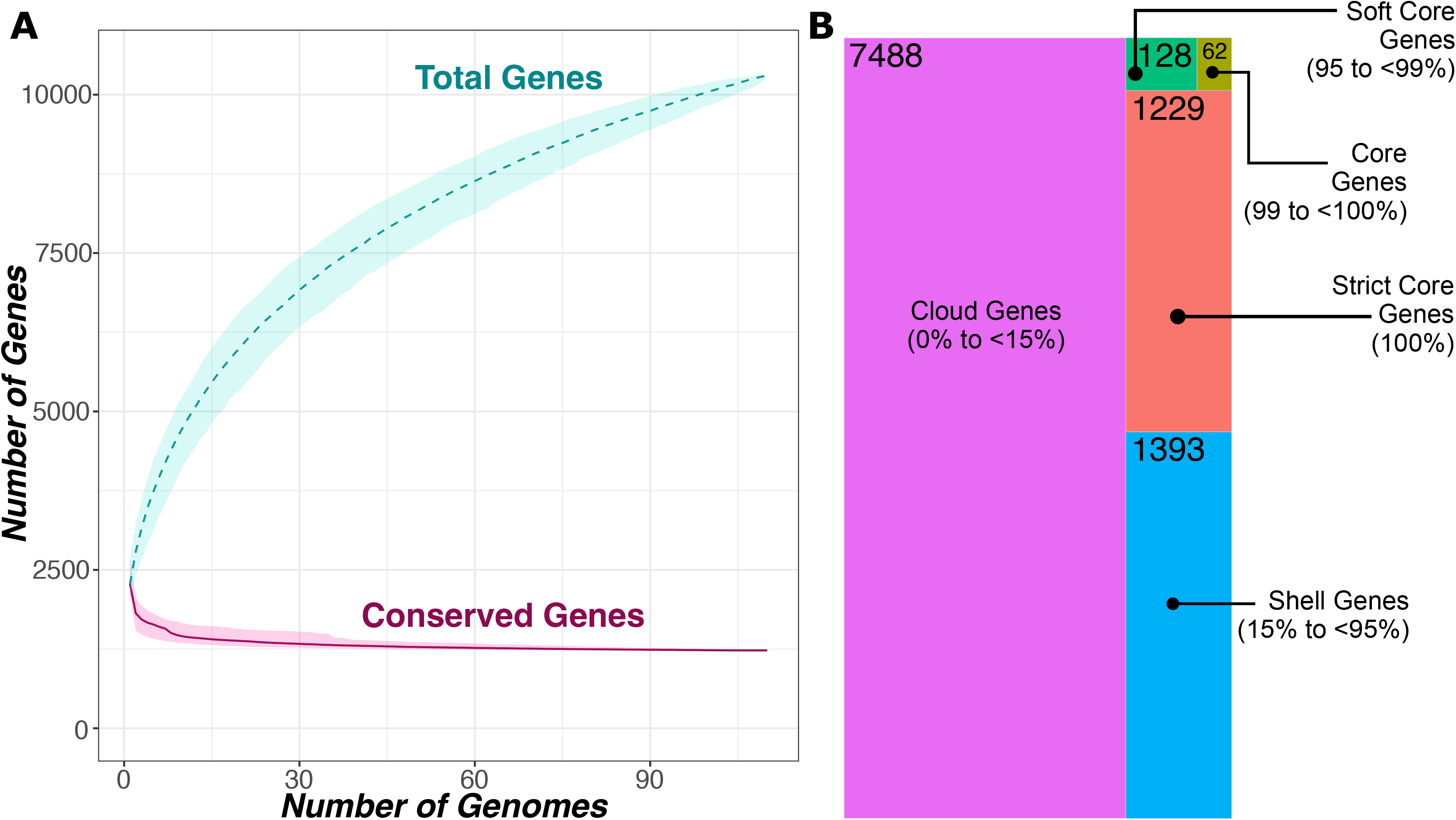
(A) Rarefaction curves for the *Macrococcus* pan- and core-genome, constructed using all 104 high-quality, publicly available *Macrococcus* genomes, plus six bovine-associated South African genomes sequenced here (*n* = 110 total *Macrococcus* genomes). Curves showcase the accumulation of pan genes (“Total Genes”) and core genes (“Conserved Genes”) using 1,000 random permutations. Dashed and solid curved lines denote median values for pan and core genes, respectively, and shading surrounding each line denotes the respective 95% confidence interval. (B) Treemap showcasing the number of genes detected within a given percentage of *Macrococcus* genomes (out of 110 total genomes). Tile sizes are proportional to the number of genes detected within a given percentage of *Macrococcus* genomes; numerical labels within each tile denote the corresponding number of genes. The treemapify v2.5.5 (https://CRAN.R-project.org/package=treemapify) R package was used to construct the plot. For both (A) and (B), PEPPAN was used to construct the core- and pan-genomes using a 40% amino acid identity threshold and a core genome threshold of 95%.

### 3.7 *Macrococcus caseolyticus* and *Macrococcus armenti* are composed of multiple within-species subclusters separated by recent gene flow

PopCOGenT identified 18 “main clusters” (species) across *Macrococcus* in its entirety; within two of these main clusters (i.e., PopCOGenT Main Clusters 0 and 2 in Figure 1), PopCOGenT identified multiple “subclusters” separated by recent gene flow (i.e., populations that were still connected by some gene flow, but had significantly more gene flow within the population than between populations; Figure 1). Specifically, (i) within PopCOGenT Main Cluster 0 (corresponding to GTDB-Tk’s *Macrococcus caseolyticus* genomospecies), five subclusters were identified, and (ii) within PopCOGenT Main Cluster 2 (an unknown species via GTDB-Tk, which contains the *M. armenti* type strain and will thus be referred to as *M. armenti* hereafter), two subclusters were identified. As such, we will discuss these two species individually in detail below (Figure 1).

#### 3.7.1 African and European *Macrococcus caseolyticus* strains largely belong to separate lineages

The 58 *Macrococcus caseolyticus* genomes (per GTDB-Tk) were divided into five PopCOGenT subclusters and five RhierBAPS clusters, although the composition of those (sub)clusters differed slightly (Figure 5, Supplementary Figure S11, and Supplementary Table S7). Notably, the majority of European *Macrococcus caseolyticus* genomes (*n* = 33 of 42 European *M. caseolyticus* genomes, 78.6%) were assigned to a well-supported clade within the species phylogeny (referred to hereafter as the “*Macrococcus caseolyticus* major European lineage”, which is denoted in Figure 5 as RhierBAPS Cluster 1; ultrafast bootstrap support = 100%). Members of the *Macrococcus caseolyticus* major European lineage were overwhelmingly of bovine origin (34 of 36 RhierBAPS Cluster 1 genomes, 94.4%), and nearly all genomes within the lineage were reportedly isolated from European countries: thirty from the United Kingdom (83.3%), and two and one genome(s) from Switzerland and Ireland, respectively (5.6% and 2.8%); the only genome reportedly isolated from outside of Europe was reportedly isolated from ropy milk in the United States in 1920 (NCBI GenBank Assembly accession GCA_900453015.1; Figure 5, Supplementary Figure S11, and Supplementary Table S3). Interestingly, the majority of genomes within the *Macrococcus caseolyticus* major European lineage were predicted to be MDR (Figures 3 and 5 and Supplementary Figure S11). Specifically, (i) all genomes in the *Macrococcus caseolyticus* major European lineage (36 of 36 genomes, 100%) were predicted to be resistant to macrolides, largely due to the presence of *abc-f* (35 of 36 *Macrococcus caseolyticus* major European lineage genomes, 97.2%; the only genome in which *abc-f* was not detected possessed *erm(B)* and was thus still predicted to be macrolide-resistant via AMRFinderPlus); (ii) nearly all (33 of 36 genomes, 91.7%) were predicted to be resistant to beta-lactams, largely due to the presence of *mecD* in 28 genomes (77.8% of 36 genomes in the lineage; the remaining five genomes that were predicted to be beta-lactam-resistant harbored *bla* and *mecB*); (iii) a majority (21 of 36 genomes in the lineage, 58.3%) were predicted to be resistant to aminoglycosides, due largely to the presence of *str* and/or *aadD1* (detected in 14 and 9 of 36 genomes, 38.9% and 25.0%, respectively; Figures 3 and 5 and Supplementary Figure S11). Additionally, three genomes possessed genes conferring resistance to bleomycin; these were the only genomes within the *Macrococcus* genus, which harbored bleomycin resistance-conferring gene *bleO* (Figures 3 and 5 and Supplementary Figure S11).

**Figure 5.**
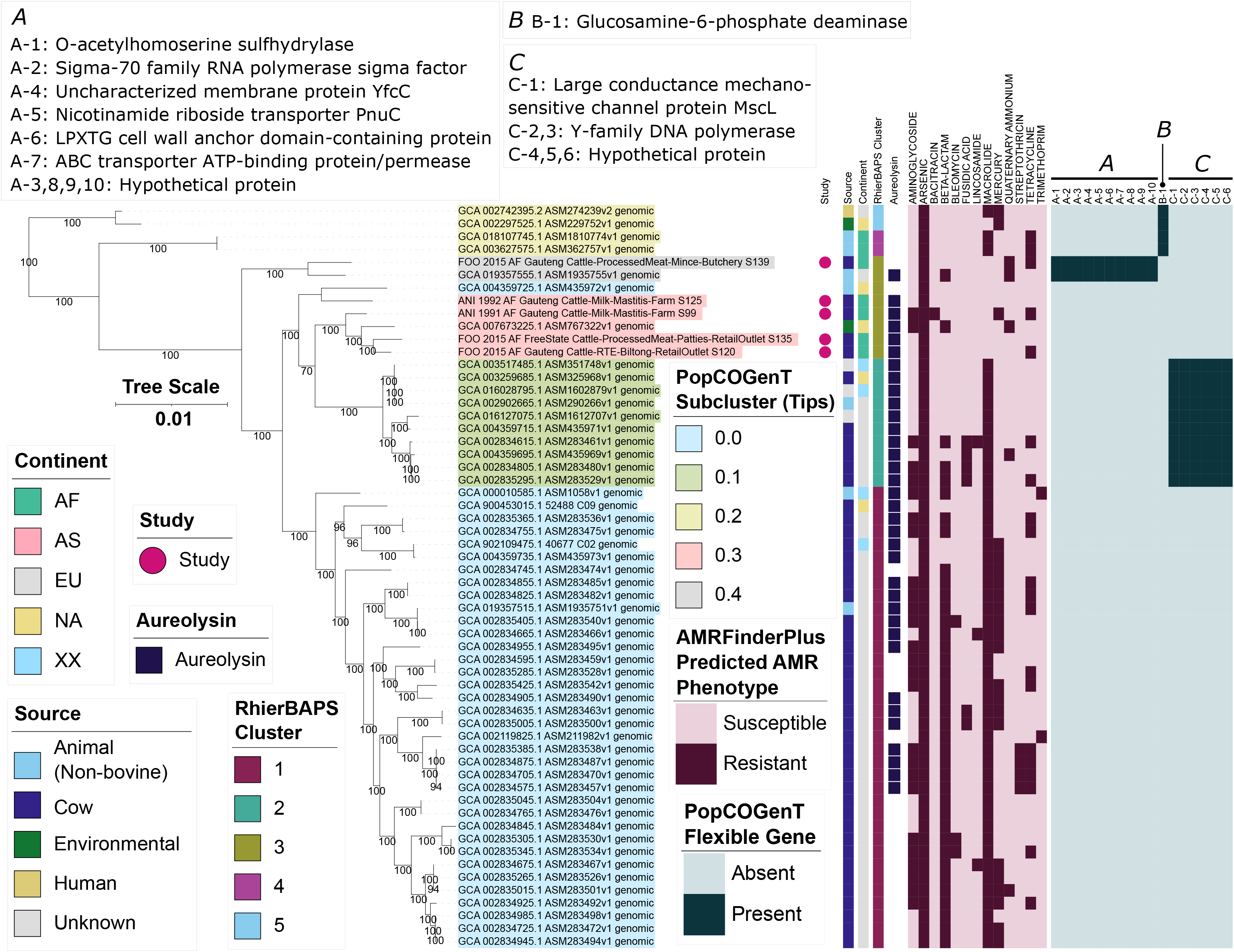
Maximum likelihood (ML) phylogeny of 58 genomes assigned to the Genome Taxonomy Database’s (GTDB) *Macrococcus caseolyticus* genomospecies. Tip label colors correspond to subcluster assignments obtained using PopCOGenT (“PopCOGenT Subcluster”). Pink circles denote genomes sequenced in this study (“Study”). Color strips/heatmaps to the right of the phylogeny denote (from left to right): (i) the source from which each strain was reportedly isolated (“Source”); (i) the continent from which each strain was reportedly isolated (“Continent”); (iii) cluster assigned using RhierBAPS (“RhierBAPS Cluster”); (iv) presence of gene(s) sharing homology to aureolysin at 40% amino acid identity and 50% coverage (“Aureolysin”); (v) predicted antimicrobial resistance (AMR) and stress response phenotype, obtained using AMR and stress response determinants identified via AMRFinderPlus (“AMRFinderPlus Predicted AMR Phenotype”); (vi) presence and absence of flexible genes identified via PopCOGenT (“PopCOGenT Flexible Gene”), with corresponding gene annotations displayed in the boxes marked “A”, “B”, and “C”. The ML phylogeny was constructed using an alignment of 1,751 core genes identified among all 58 *Macrococcus caseolyticus* genomes, plus an outgroup *Macrococcus* spp. genome from bactaxR Cluster 2 (NCBI GenBank Assembly accession GCA_019357535.1; Figure 1), using Panaroo and a 70% protein family sequence identity threshold. The tree was rooted using the outgroup (omitted for readability), and branch lengths are reported in substitutions per site. Branch labels correspond to branch support percentages obtained using one thousand replicates of the ultrafast bootstrap approximation. AF, Africa; AS, Asia; EU, Europe; NA, North America; XX, unknown/unreported geographic location. For an extended version of this phylogeny, see Supplementary Figure S11.

Of the nine European *Macrococcus caseolyticus* genomes that were not members of the *Macrococcus caseolyticus* major European lineage, seven belonged to a well-supported clade containing ten genomes (ultrafast bootstrap support = 100%; referred to hereafter as the “*Macrococcus caseolyticus* minor European lineage”, which is denoted in Figure 5 as RhierBAPS Cluster 2 and PopCOGenT Subcluster 0.1). Aside from two genomes of unknown origin, one genome within the *Macrococcus caseolyticus* minor European lineage was reportedly of non-European origin (i.e., strain CCM 3540, reportedly isolated from cow’s milk in the Washington, D.C. vicinity of the United States in 1916; NCBI GenBank Assembly accession GCA_003259685.1, Supplementary Table S3) (Evans, 1916). Like the *Macrococcus caseolyticus* major European lineage, all genomes within the *Macrococcus caseolyticus* minor European lineage were predicted to be resistant to macrolides, as all harbored *abc-f* (Figures 3 and 5 and Supplementary Figure S11). However, a predicted MDR phenotype (i.e., resistant to three or more antimicrobial classes) was less prevalent among genomes within the minor European lineage (*n* = 3 of 10 *Macrococcus caseolyticus* minor European lineage genomes, 30%): the MDR genomes were similar on a genomic scale (99.7-99.9 ANI via OrthoANI) and were confined to a single, well-supported clade within the *Macrococcus caseolyticus* minor European lineage (ultrafast bootstrap support = 100%; Figure 5 and Supplementary Figure S11). Additionally, within the *Macrococcus caseolyticus* minor European lineage, PopCOGenT identified six “flexible” genes (i.e., PopCOGenT subcluster-specific orthologous gene clusters), which were specific to the *Macrococcus caseolyticus* minor European lineage (denoted as gene group “C” within the PopCOGenT Flexible Gene heatmap in Figure 5; PopCOGenT *P* < 0.05). All six genes were chromosomal and included (i) large conductance mechanosensitive channel protein MscL, and (ii) genes associated with Y-family DNA polymerases (Figure 5, Supplementary Figure S11, and Supplementary Table S8). Compared to all other *Macrococcus caseolyticus* genes, numerous biological processes (BPs) and molecular functions (MFs) were enriched in the *Macrococcus caseolyticus* minor European lineage flexible genes, including DNA-related BPs/MFs (e.g., DNA biosynthesis, replication, and repair), and those related to ion binding/transport (topGO Fisher’s Exact Test [FET] *P* < 0.05; Supplementary Tables S8 and S19).

Of the 12 *Macrococcus caseolyticus* genomes, which were not members of the major and minor European lineages, seven were African in origin, three were North American, and two were European, including the one human-associated *Macrococcus caseolyticus* genome (i.e., strain CCM 7927, which was isolated in Pribram, Czech Republic in 2003 from a vaginal swab taken from an acute vaginitis case in a 40-45 year-old patient, NCBI GenBank Assembly accession GCA 002742395.2; Figure 5, Supplementary Figure S11, and Supplementary Table S3) (Maslanova et al., 2018). Notably, of the five South African *Macrococcus caseolyticus* strains isolated and sequenced here, four were assigned to a single PopCOGenT subcluster (i.e., PopCOGenT Subcluster 0.3 in Figure 5). Unlike the major and minor European lineages, members of this subcluster did not possess macrolide resistance genes (Figures 3 and 5 and Supplementary Figure S11). AMR genes were detected sporadically within these genomes. Specifically, (i) strain S99 possessed genes associated with aminoglycoside (streptomycin), bacitracin, and tetracycline resistance (*str*, *bcrBC*, and *tet(L)*, respectively); (ii) GCA_007673225.1 (an environmental strain isolated in 2018 in Durham, North Carolina, United States) possessed genes associated with aminoglycoside and beta-lactam resistance (i.e., *aph(2’’)-IIIa*, *str*, and *mecD*, associated with amikacin/gentamicin/kanamycin/tobramycin, streptomycin, and methicillin resistance, respectively); (iii) S120 possessed tetracycline resistance gene *tet(L)* (Figure 5 and Supplementary Figure S11). Despite most genomes being South African in origin, the five *Macrococcus caseolyticus* genomes within this subcluster were considerably diverse, sharing 98.6-99.4 ANI with each other (via OrthoANI; Figure 5 and Supplementary Figure S11).

The remaining South African genome sequenced in this study (i.e., S139), plus GCA_019357555.1 (isolated from a calf nasal swab in Switzerland in 2019), were assigned to a separate subcluster via PopCOGenT (i.e., PopCOGenT Subcluster 0.4 in Figure 5). Neither genome possessed macrolide resistance genes, although both possessed quaternary ammonium resistance gene *qacH* (Figure 5 and Supplementary Figure S11). S139 additionally possessed tetracycline resistance gene *tet(L)*, while GCA_019357555.1 possessed beta-lactam resistance genes *mecB* (methicillin) and *bla* (Figure 5 and Supplementary Figure S11). Most notably, however, PopCOGenT identified ten flexible genes within this subcluster (denoted as gene group “A” within the PopCOGenT Flexible Gene heatmap in Figure 5, PopCOGenT *P* < 0.05; Figure 5, Supplementary Figure S11, and Supplementary Table S8); ATP- and transmembrane-associated BPs/MFs were enriched in this subcluster’s flexible genes (topGO FET *P* < 0.05; Supplementary Table S21).

Four additional *Macrococcus caseolyticus* genomes were assigned to a single subcluster using PopCOGenT (i.e., PopCOGenT Subcluster 0.2 in Figure 5). Interestingly, like the major and minor European clades, three of the four genomes within this subcluster were predicted to be macrolide resistant, as they possessed *abc-f* and *mef(D)* (Figure 5 and Supplementary Figure S11). Two highly similar genomes derived from strains isolated in 2016 from wounded animals in Sudan additionally possessed tetracycline resistance gene *tet(L)* (100.0 ANI and 0 SNPs via OrthoANI and Snippy, respectively, NCBI GenBank Assembly accessions GCA_018107745.1 and GCA_003627575.1; Figure 5 and Supplementary Figure S11). Additionally, unlike the other *Macrococcus caseolyticus* subclusters described above, none of the genomes within this PopCOGenT subcluster possessed genes sharing homology to aureolysin-encoding genes (Figure 5 and Supplementary Figure S11). Further, PopCOGenT identified one flexible gene within this subcluster (denoted as gene group “B” within the PopCOGenT Flexible Gene heatmap in Figure 5, PopCOGenT *P* < 0.05; Supplementary Table S8): glucosamine-6-phosphate deaminase, which was associated with the enrichment of several GO terms, including antibiotic catabolic process, carbohydrate metabolic process, and N-acetylglucosamine-associated processes (topGO FET *P* < 0.05; Figure 5, Supplementary Figure S11, and Supplementary Tables S8 and S20).

Overall, these results indicate that *Macrococcus caseolyticus* genomes from geographic regions outside of Europe, particularly Africa, belong to separate lineages within the species. However, future genomic sequencing efforts are needed to provide further evidence of lineage-geography associations.

#### 3.7.2 Putative virulence factors are differentially associated with *Macrococcus armenti* lineages

Like *Macrococcus caseolyticus*, *Macrococcus armenti* could be differentiated into subclusters via PopCOGenT (Figure 6A, Supplementary Figure S12, and Supplementary Table S7). Specifically, PopCOGenT Subcluster 2.0 contained five genomes from animals in Switzerland (two from strains isolated from the nasal cavities of calves in 2019, and three from the skins of pigs in 2021); and (ii) PopCOGenT Subcluster 2.1 contained two genomes from pigs in Switzerland (one from the nasal cavity of a pig in 2017, and another from the skin of a pig in 2021; Figure 6A, Supplementary Figure S12, and Supplementary Tables S3 and S7). An additional genome, derived from a pig-associated strain isolated in the United Kingdom in 1963 (NCBI GenBank Assembly accession GCA_022808015.1) was additionally assigned to the *Macrococcus armenti* species via ANI-based methods (i.e., using OrthoANI, it shared 96.5-97.7 ANI with all other *Macrococcus armenti* genomes; Figure 2); however, PopCOGenT assigned this genome to a separate main cluster (i.e., “species”), and it was thus not included in the subsequent within-main cluster flexible gene analyses (Figure 6A, Supplementary Figure S12, and Supplementary Table S7).

**Figure 6.**
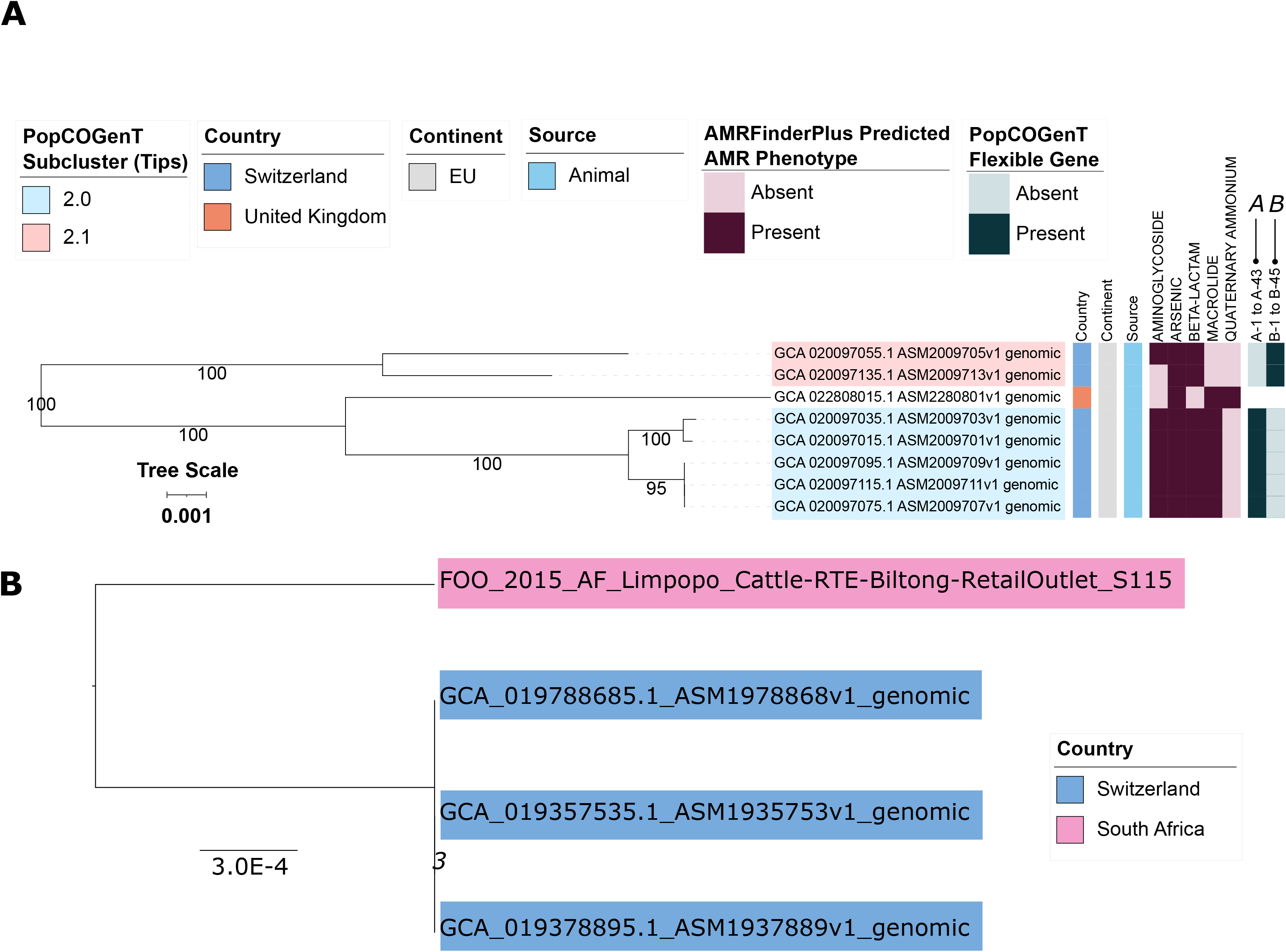
(A) Maximum likelihood (ML) phylogeny of eight genomes assigned to bactaxR Cluster 13 (i.e., *Macrococcus armenti*, based on average nucleotide identity [ANI]-based comparisons to species type strain genomes; Figure 1). Tip label colors correspond to subcluster assignments obtained using PopCOGenT (“PopCOGenT Subcluster”; one genome was not assigned to the same main cluster via PopCOGenT, and thus is not colored). Color strips/heatmaps to the right of the phylogeny denote (from left to right): (i) the country from which each strain was reportedly isolated (“Country”); (ii) the continent from which each strain was reportedly isolated (“Continent”); (iii) the source from which each strain was reportedly isolated (“Source”); (iv) predicted antimicrobial resistance (AMR) and stress response phenotype, obtained using AMR and stress response determinants identified via AMRFinderPlus (“AMRFinderPlus Predicted AMR Phenotype”); (v) presence and absence of flexible genes identified via PopCOGenT (“PopCOGenT Flexible Gene”; for gene descriptions, see Supplementary Table S9). The ML phylogeny was constructed using an alignment of 1,416 core genes identified among all eight *Macrococcus armenti* genomes, plus an outgroup *Macrococcus canis* genome (NCBI GenBank Assembly accession GCA_014524485.1; Figure 1), using Panaroo and a 70% protein family sequence identity threshold. The tree was rooted using the outgroup (omitted for readability), and branch lengths are reported in substitutions per site. Branch labels correspond to branch support percentages obtained using one thousand replicates of the ultrafast bootstrap approximation. EU, Europe. For an extended version of this phylogeny, see Supplementary Figure S12. (B) ML phylogeny constructed using core SNPs identified among four genomes assigned to bactaxR Cluster 2, a putative novel GTDB genomospecies, which shares >95 ANI with several *Macrococcus caseolyticus* genomes but < 95 ANI with others (Figures 1 and 2). Tip label colors correspond to reported country of isolation. Core SNPs were identified using Snippy, filtered using Gubbins/snp-sites, and the phylogeny was constructed using IQ-TREE. The phylogeny is rooted at the midpoint, and branch lengths are reported in substitutions per site. Branch labels correspond to branch support percentages obtained using one thousand replicates of the ultrafast bootstrap approximation.

Within PopCOGenT Subcluster 2.0, PopCOGenT identified 43 flexible genes (denoted as gene group “A” within the PopCOGenT Flexible Gene heatmap in Figure 6A, PopCOGenT *P* < 0.05; Supplementary Table S9), which together were associated with the enrichment of eight GO terms (topGO FET *P* < 0.05; Supplementary Table S22). The most highly enriched GO terms were by far “diaminopimelate biosynthetic process” (GO:0019877) and “lysine biosynthetic process via diaminopimelate” (GO:0009089, topGO FET *P* < 1.0×10^-30^; Supplementary Table S22), which were assigned to a cluster of three consecutive flexible genes (PopCOGenT *P* < 0.05): (i) 4-hydroxy-tetrahydrodipicolinate reductase/dihydrodipicolinate reductase *dapB* (NCBI Protein accession UBH07557.1); (ii) 2,3,4,5-tetrahydropyridine-2,6-dicarboxylate N-acetyltransferase *dapD* (NCBI Protein accession UBH09720.1); (iii) an amidohydrolase (NCBI Protein accession UBH07558.1; Supplementary Table S9).

Most notably, genes sharing homology to *Staphylococcus aureus* Type VII secretion system proteins were among the flexible genes within Subcluster 2.0 (PopCOGenT *P* < 0.05), including genes sharing homology to extracellular protein EsxD (VFDB ID VFG049714), chaperone protein EsaE (VFDB ID VFG049701), secreted protein EsxB (VFDB ID VFG002411), secretion substrate EsaC (at 97% query coverage and 38% AA identity; NCBI Protein accession HCD1544785.1), EssB (NCBI Protein accession UBH08107.1), EsaA (NCBI Protein accession UBH08110.1), and secreted protein EsxA (VFDB ID VFG002405; Figure 6A, Supplementary Figure S12, and Supplementary Table S9).

Several additional clusters of genes were among the flexible genes within Subcluster 2.0 (PopCOGenT *P* < 0.05; Figure 6A, Supplementary Figure S12, and Supplementary Table S9), including: (i) a cluster of genes involved in nitrous oxide reduction, e.g., c-type cytochrome (NCBI Protein accession UBH08788.1), a Sec-dependent nitrous-oxide reductase (NCBI Protein accession UBH08789.1), nitrous oxide reductase family maturation protein NosD (NCBI Protein accession UBH08791.1); (ii) a cluster of genes that included an ImmA/IrrE family metallo-endopeptidase (NCBI Protein accession UBH09010.1), a LacI family DNA-binding transcriptional regulator (NCBI Protein accession UBH09033.1), a sucrose-6-phosphate hydrolase (NCBI Protein accession UBH09034.1), a carbohydrate kinase (NCBI Protein accession UBH09035.1), and sucrose-specific PTS transporter subunit IIBC (NCBI Protein accession UBH09036.1); (iii) a cluster of genes that included a pathogenicity island protein (NCBI Protein accession UBH09209.1; Figure 6A, Supplementary Figure S12, and Supplementary Table S9).

Interestingly, a protein most closely resembling immune inhibitor A was also identified by PopCOGenT as a flexible gene (at 98% query coverage and 97.65% AA identity, NCBI Protein accession WP_224185801.1, PopCOGenT *P* < 0.05; Figure 6A, Supplementary Figure S12, and Supplementary Table S9). The “immune inhibitor A peptidase M6” protein domain identified in this protein (PFAM ID 05547) has previously been identified in virulence factors secreted by members of the *Bacillus cereus* group (immune inhibitor A; InhA) and *Vibrio cholerae* (secreted metalloprotease PrtV) (Vaitkevicius et al., 2008).

Comparatively, within Subcluster 2.1, PopCOGenT identified 45 flexible genes (denoted as gene group “B” within the PopCOGenT Flexible Gene heatmap in Figure 6A, PopCOGenT *P* < 0.05) associated with 22 enriched GO terms (topGO FET *P* < 0.05; Figure 6A, Supplementary Figure S12, and Supplementary Tables S9 and S23). By far the most highly enriched GO term within this subcluster corresponded to BP “lipoteichoic acid biosynthetic process” (GO:0070395, topGO FET *P* = 2.2×10^-18^; Supplementary Table S23). Notably, a cluster of five consecutive, chromosomally encoded flexible genes within PopCOGenT Subcluster 2.1 were associated with (lipo)teichoic acid synthesis (PopCOGenT *P* < 0.05; Supplementary Table S9): teichoic acid D-Ala incorporation-associated protein DltX (NCBI Protein accession UBH13741.1), D-alanine--poly(phosphoribitol) ligase subunit DltA (NCBI Protein accession UBH13742.1), D-alanyl-lipoteichoic acid biosynthesis protein DltB (NCBI Protein accession UBH13743.1), D-alanine--poly(phosphoribitol) ligase subunit 2 DltC (NCBI Protein accession UBH13744.1), and D-alanyl-lipoteichoic acid biosynthesis protein DltD (NCBI Protein accession UBH13745.1). Interestingly, this cluster of five genes was located several genes downstream of two consecutive, chromosomally encoded beta-lactamase family proteins, which were also both identified as being flexible genes (PopCOGenT *P* < 0.05). Both beta-lactamase family proteins were annotated via eggNOG-mapper as “autolysis and methicillin resistant-related protein PbpX” (NCBI Protein accessions UBH13736.1 and UBH13737.1) and were associated with “response to antibiotic” (GO:0046677), a BP that was also enriched in PopCOGenT Subcluster 2.1 (topGO FET *P* = 2.3×10^-3^; Supplementary Tables S9 and S23).

Several GO terms associated with transporter activity were also enriched in PopCOGenT Subcluster 2.1 (topGO FET *P* < 0.05), including MF “ABC−type transporter activity” (GO:0140359; Supplementary Table S23). Congruently, four separate clusters of genes containing regions annotated as ABC transporter components were included among PopCOGenT’s set of flexible genes (PopCOGenT *P* < 0.05; Supplementary Tables S9 and S23)

Interestingly, a protein annotated as immune inhibitor A was also among the flexible genes detected within PopCOGenT Subcluster 2.1 (PopCOGenT *P* < 0.05, NCBI Protein accession UBH13622.1; Figure 6A, Supplementary Figure S12, and Supplementary Table S9). Further, genes encoding a type II toxin-antitoxin system were among the flexible genes identified by PopCOGenT within this PopCOGenT subcluster (PopCOGenT *P* < 0.05; Figure 6A, Supplementary Figure S12, and Supplementary Table S9), specifically: (i) a type II toxin-antitoxin system RelE/ParE family toxin, which was immediately upstream of (ii) a type II toxin-antitoxin system Phd/YefM family antitoxin (NCBI Protein accessions UBH12746.1 and UBH12747.1, respectively).

Overall, (i) *Macrococcus armenti* boasts two major subclusters, which are largely separated by recent gene flow; and (ii) flexible genes differentially present within these major subclusters (e.g., a type VII secretion system, toxin-antitoxin genes, beta-lactamase family genes) indicate that these two subclusters may differ phenotypically, although future experiments will be necessary to confirm this.

### 3.8 A novel GTDB genomospecies encompasses *Macrococcus* genomes from Switzerland and South Africa

Of the six *Macrococcus* spp. genomes sequenced in this study, five were assigned to *M. caseolyticus* (per GTDB-Tk; Figure 1 and Supplementary Table S4). The genome of S115, however, could not be assigned to a known species via GTDB-Tk (Figure 1 and Supplementary Table S4). Using bactaxR and a 95 ANI threshold (i.e., an approach similar to that of GTDB-Tk), three additional, publicly available genomes belonged to this putative novel GTDB genomospecies (i.e., bactaxR Cluster 2, *n* = 4 total genomes; Figures 1 and 6B). In addition to (i) S115, a food-associated strain isolated in 2015 from beef biltong sold in South Africa’s Limpopo province, this genomospecies included three strains isolated in Switzerland in 2019: (ii) 19Msa1099, isolated from pork meat (NCBI GenBank Assembly accession GCA_019357535.1), plus (iii) 19Msa1047 and (iv) 19Msa0499, each isolated from calf nasal swab samples (NCBI GenBank Assembly accessions GCA_019378895.1 and GCA_019788685.1, respectively; Supplementary Tables S1 and S3).

Notably, the South African genome was relatively distantly related to the Swiss genomes, sharing 99.2 ANI with each via OrthoANI and differing by 8,614-8,637 SNPs (identified via Snippy relative to each individual Swiss genome).

Comparatively, the three Swiss genomes shared >99.99 ANI with each other via OrthoANI and differed by 1-34 core SNPs (calculated via Snippy with the South African S115 strain excluded): strains 19Msa1047 (from a calf nasal swab) and 19Msa1099 (from pork meat) differed by a single core SNP identified in a gene annotated as a CBS domain-containing protein (NCBI Protein accession WP_219491817.1, corresponding to locus tag KYI07_RS05750 within the *M. caseolyticus* str. 19Msa0499 reference chromosome with NCBI Nucleotide accession NZ_CP079969.1). These two strains differed from strain 19Msa0499 (from a calf nasal swab) by 33 and 34 core SNPs, all of which fell within two regions of the *M. caseolyticus* str. 19Msa0499 reference chromosome: (i) 13 core SNPs within positions 312,553-367,746 bp, and (ii) 20 core SNPs within positions 1,778,236-1,778,444 bp, indicating that genetic differences within these regions may be due to recombination.

## 4 Discussion

In this study, WGS was used to characterize *Macrococcus* spp. strains isolated from South African cattle (i.e., two strains from bovine clinical mastitis cases) and beef products (i.e., two stains from RTE beef biltong and two from minced/processed beef products). Using these genomes in combination with all publicly available, high quality *Macrococcus* spp. genomes, insight is provided into the evolution, population structure, and functional potential of the *Macrococcus* genus as a whole. Importantly, we observed (i) differences in functional potential (e.g., AMR potential, virulence potential) between and within *Macrococcus* spp., and (ii) that some *Macrococcus* species lack clear boundaries at conventional genomospecies delineation thresholds (i.e., 95 ANI), which may cause taxonomic issues in the future. Below, we discuss these findings in detail, as well as (iii) future opportunities in the *Macrococcus* genomics space.

### 4.1 Differences in functional potential can be observed between and within *Macrococcus* species

Bacteria can adapt to stressors and stimuli in their respective environments through the acquisition of genomic material in the “flexible” gene pool (Arevalo et al., 2019). Thus, intraspecies differences in genomic content can be observed for many bacterial species (Tonkin-Hill et al., 2020), and differences resulting from recent gene flow (i.e., genomic elements acquired post-speciation) can be used to delineate populations within those species (Arevalo et al., 2019). Here, we queried all *Macrococcus* spp. genomes and identified genomic determinants variably present within species, indicative of within-species differences in functional potential. For example, of the 50 putative AMR and stress response determinants identified across *Macrococcus* in its entirety, nearly half (24 of 50, 48%) were species-specific (based on GTDB-Tk species assignments); of these species-specific AMR and stress response determinants, all (24 of 24, 100%) were variably present within their given species, indicating that AMR potential can vary within *Macrococcus* species. Antimicrobial exposure can select for AMR (Hendriksen et al., 2019; Olesen et al., 2020), and reducing exposure (e.g., limiting antimicrobial use outside of treating human disease, minimizing unnecessary antibiotic use for human illness cases) can reduce the risk of AMR (Antimicrobial Resistance Collaborators, 2022). Thus, it is not particularly surprising that intraspecies differences in AMR potential exist within *Macrococcus*; the genomes aggregated here were derived from *Macrococcus* strains isolated from a range of sources (e.g., humans, animal hosts, animal products, environmental samples), geographic locations (i.e., four continents), and timeframes (i.e., between the years of 1916 and 2021) and thus have likely been exposed to different selective pressures.

Comparatively, some genomic elements identified here were present across multiple *Macrococcus* spp., indicating shared inter-species functional potential for some phenotypes. Methicillin resistance genes *mecB* and *mecD*, for example, were variably present within multiple *Macrococcus* species (via GTDB-Tk; Figure 3), mirroring previous studies, which have reported *mecB* and/or *mecD* in various *Macrococcus* spp., including *M. caseolyticus* (Schwendener et al., 2017; MacFadyen et al., 2018; Zhang et al., 2022), *M. bohemicus* (Foster and Paterson, 2020), *M. canis* (Chanchaithong et al., 2019), and *M. goetzii* (Maslanova et al., 2018). Outside of the AMR space, we further identified proteins that shared homology with virulence factors in other species. Perhaps most notably, we detected homologues of aureolysin in multiple *Macrococcus* species (Figure 3). Aureolysin is an extracellular zinc-dependent metalloprotease secreted by *Staphylococcus aureus*, which plays a crucial role in host immune system evasion (Thammavongsa et al., 2015; Pietrocola et al., 2017). While others have detected aureolysin homologues in *Macrococcus* genomes (Mazhar et al., 2019a; b; Zhang et al., 2022), the roles this protein plays in *Macrococcus* interactions with human or animal hosts (if any) are unknown.

Finally, for *Macrococcus caseolyticus* and *Macrococcus armenti*, which were composed of multiple populations (subclusters) separated by recent gene flow, some variably present genomic elements were subcluster-specific genes, which had been acquired post-speciation and differentially swept through these subclusters (i.e., flexible genes identified via PopCOGenT). Similar to results observed for *Ruminococcus gnavus* (Arevalo et al., 2019), transporter functions (e.g., ABC-type transporters, genes involved in ion transport) were enriched in subcluster-specific flexible gene sets within both *M. caseolyticus* and *M. armenti*. Notably, within *M. armenti*, we further identified two distinct subclusters with different flexible genes in each, including (i) one subcluster with a type VII secretion system, *Staphylococcus aureus*-like virulence factors, and a putative pathogenicity island (Subcluster 2.0), and (ii) another with beta-lactamase family proteins and a type II toxin-antitoxin system (Subcluster 2.1). Taken together, these results indicate that there may be differences in the functional potential of these two *M. armenti* subclusters; however future experimental work will be needed to confirm the roles of these subcluster-specific flexible genes in each subcluster, as there are no clear differences in terms of each subcluster’s ecological niche (strains in both subclusters were isolated from livestock in Switzerland).

Overall, proteins with potential virulence- and AMR-related functions, which were differentially present within and across *Macrococcus* species were identified. This indicates that there are potential within- and between-species differences in *Macrococcus* virulence and AMR potential. Future experimental efforts will thus be needed to investigate these differences further.

### 4.2 The lack of clear genomospecies boundaries between some *Macrococcus* species may cause taxonomic issues in the future

The delineation of prokaryotes into species-level taxonomic units is notoriously challenging, as horizontal gene transfer can obscure prokaryotic population boundaries (Jain et al., 2018; Arevalo et al., 2019). With the increasing availability of WGS, taxonomic assignment has largely shifted to *in silico* methods; however, numerous approaches exist for this purpose and may produce conflicting results (e.g., various implementations of ANI-based methods, marker gene-based methods, metrics using recent gene flow, *in silico* DNA-DNA hybridization) (Meier-Kolthoff et al., 2013; Mende et al., 2013; Lee et al., 2016; Yoon et al., 2017; Jain et al., 2018; Arevalo et al., 2019; Chaumeil et al., 2019; Meier-Kolthoff et al., 2022; Parks et al., 2022). Here, we applied multiple species-level taxonomic assignment methods to all publicly available *Macrococcus* genomes, specifically ANI-based approaches (i.e., OrthoANI/bactaxR and GTDB-Tk), an approach that uses a metric based on recent gene flow (i.e., PopCOGenT), and a marker gene-based method (i.e., specI; Figure 1). Overall, we observed similar results for three of four approaches; the marker gene-based approach only recovered two species, likely due to a lack of *Macrococcus* genomes of species other than *M. caseolyticus* and *M. canis* during species cluster database construction (this will likely be remedied in future specI database versions). However, even among the approaches that produced highly similar results, no two methods produced identical results.

Furthermore, at the conventional 95 ANI genomospecies threshold, several *Macrococcus* species were found to overlap (i.e., members of one species shared ≥95 ANI with members of a different species; Figure 2). We have previously observed a similar phenomenon among members of the *Bacillus cereus* group (Carroll et al., 2020), as others have done for *Escherichia/Shigella* spp., *Mycobacterium* spp., and *Neisseria gonorrhoeae/Neisseria meningitidis* (Jain et al., 2018). For *Macrococcus*, ambiguous species boundaries may not seem immediately concerning, as members of the genus are often viewed as animal commensals (Mazhar et al., 2018); thus, taxonomic misidentifications may not be viewed as “high consequence” compared to other organisms plagued by taxonomic issues (e.g. anthrax-causing organisms within the *Bacillus cereus* group, botulinum neurotoxin-producing members of *Clostridium*) (Smith et al., 2018; Bower et al., 2022). However, as more *Macrococcus* strains undergo WGS and more is learned about the pathogenic potential of these organisms in animals and humans, there may be a greater need to ensure that species are clearly defined (e.g., in clinical laboratory, diagnostic, or regulatory settings). While there is some evidence that one of the South African genomes sequenced here belongs to a putative novel species (i.e., S115), we do not advocate for any changes to the taxonomy at this time, due to the limited number of genomes available. However, we encourage readers to be aware of ambiguous species boundaries for some *Macrococcus* spp., which may cause taxonomic issues in the future.

### 4.3 Future genomic sequencing, metadata collection, and phenotypic characterization efforts are needed to gain insight into *Macrococcus* population structure, antimicrobial resistance, and virulence potential

WGS has proven to be revolutionary in the food, veterinary, and human clinical microbiology spaces and is being used for—among other applications—pathogen surveillance, outbreak and cluster detection, source tracking, and diagnostics (Rossen et al., 2018; Brown et al., 2021; Ferdinand et al., 2021; Forde et al., 2022). Massive WGS efforts are being undertaken to query bacterial pathogens such as *Salmonella enterica*, *Escherichia coli*, and *Listeria monocytogenes* (Allard et al., 2016; Stevens et al., 2017; Brown et al., 2019), and large amounts of genomic data and metadata are publicly available for these organisms. As of 5 February 2023, (i) 455,330 genomes had been submitted to NCBI’s GenBank Assembly database as *Salmonella enterica*, (ii) 200,204 as *Escherichia coli*, and (iii) 51,579 as *Listeria monocytogenes*. *Staphylococcus aureus* is a close relative of *Macrococcus* and boasts a total of 68,631 publicly available, assembled genomes (per NCBI’s GenBank Assembly database, accessed 5 February 2023). These numbers dwarf those of *Macrococcus,* with 110 available, high-quality genomes for the entire genus at the time of this study, including the genomes generated here.

While our study provides insight into the evolution, population structure, and functional potential of *Macrococcus*, much more needs to be done to understand the role that *Macrococcus* spp. play as animal commensals, in animal-associated foodstuffs, and as opportunistic pathogens in animals and humans. First and foremost, future WGS efforts are needed to characterize these organisms, as increased availability of genomes will provide further insight into *Macrococcus* evolution (e.g., facilitating the identification of novel species, novel lineages within species). It is equally important that future WGS efforts are complemented with publicly available metadata (e.g., information conveying when and where a given strain was isolated) as this information can be used to identify potential host or geographic associations or potential migration or transmission events (e.g., between hosts or geographic regions). Finally, phenotypic data will be essential to confirm or invalidate the preliminary findings posited here regarding *Macrococcus* functional potential. Genomic AMR prediction, for example, does not necessarily translate to phenotypic AMR (Ransom et al., 2020). Similarly, any genomic determinants identified here based on their homology to known virulence factors (e.g., aureolysin, PVL, immune inhibitor A, the type VII secretion system identified in one *M. armenti* subcluster) must be evaluated experimentally. Thus, we hope that the results provided here can serve as a guide for further studies of the AMR and virulence potential of *Macrococcus* spp.

## 6 Author Contributions

LMC performed all computational analyses. IM performed bacterial isolation and identification as well as DNA extraction. RP supervised the sequencing of the isolates. IM and KM sourced the funding for sequencing of the isolates. All authors contributed to the article and approved the submitted version.

## 8 Data Availability Statement

Genomes sequenced in this study have been deposited in NCBI under BioProject accession PRJNA941163, with NCBI BioSample accession numbers for individual strains available in Supplementary Table S1. NCBI BioSample and Assembly accession numbers for all publicly available genomes used in this study are available in Supplementary Table S3.

