## Supplementary Figure S2 for "Genus-Wide Genomic Characterization of *Macrococcus*: Insights into Evolution, Population Structure, and Functional Potential"

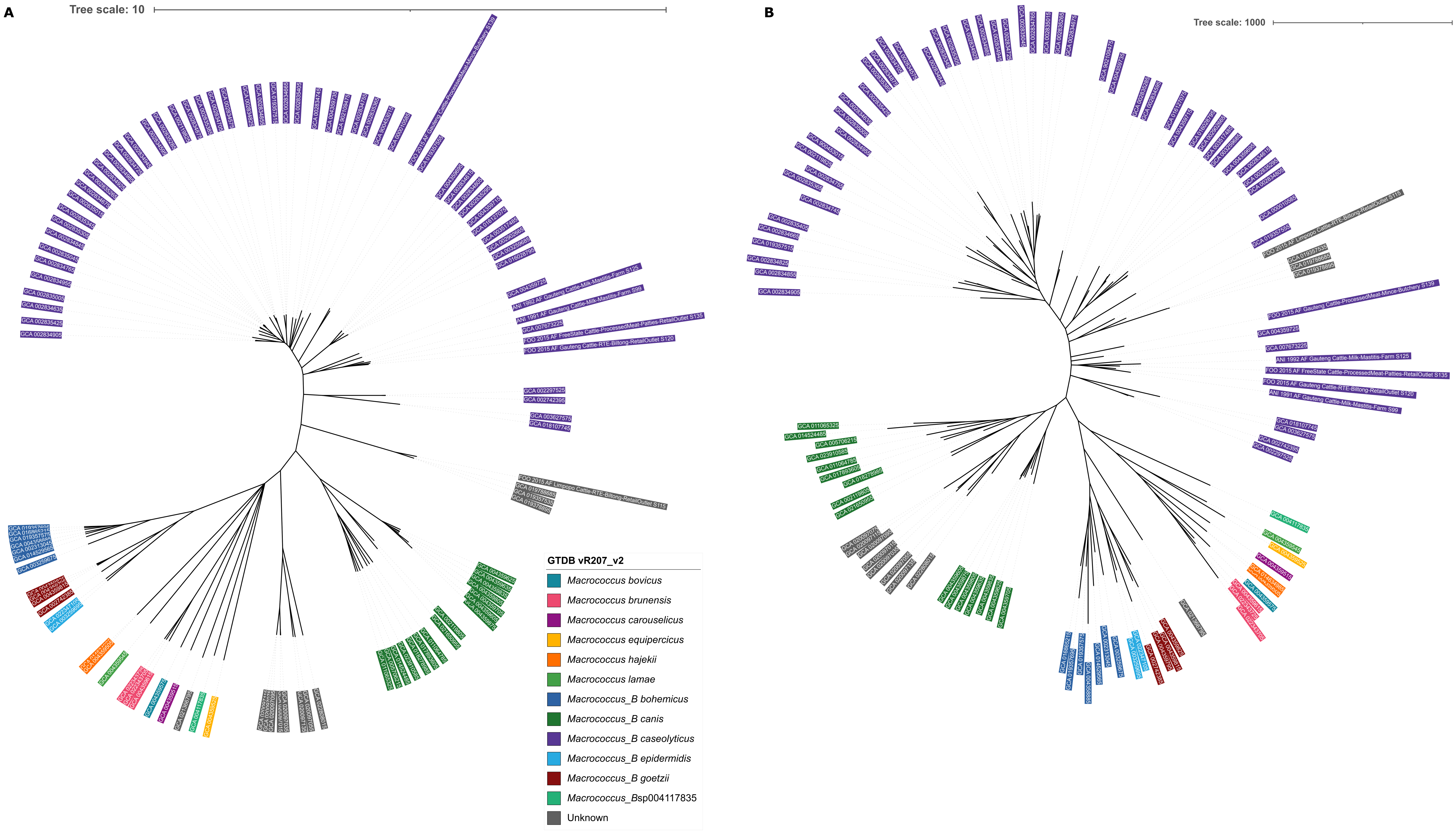


Supplementary Figure S2. Unrooted (A) Core Genome Allelic Variation (CGAV) tree and (B) accessory gene presence/absence tree, constructed using PEPPAN, a 20% amino acid identity threshold, and a 95% core genome threshold (n = 110 Macrococcus genomes). Tip label colors correspond to Genome Taxonomy Database (GTDB) species, assigned using the Genome Taxonomy Database Toolkit (GTDB-Tk) v2.1.0 and GTDB vR207_v2. PEPPAN constructed the (A) CGAV tree using RapidNJ based on numbers of identical sequences (i.e., alleles) of single copy genes present in ≥95% of Macrococcus genomes. For (B), PEPPAN used FastTree to construct the tree, using the binary presence/absence of accessory genes.
