## Supplementary Figure S3 for "Genus-Wide Genomic Characterization of *Macrococcus*: Insights into Evolution, Population Structure, and Functional Potential"

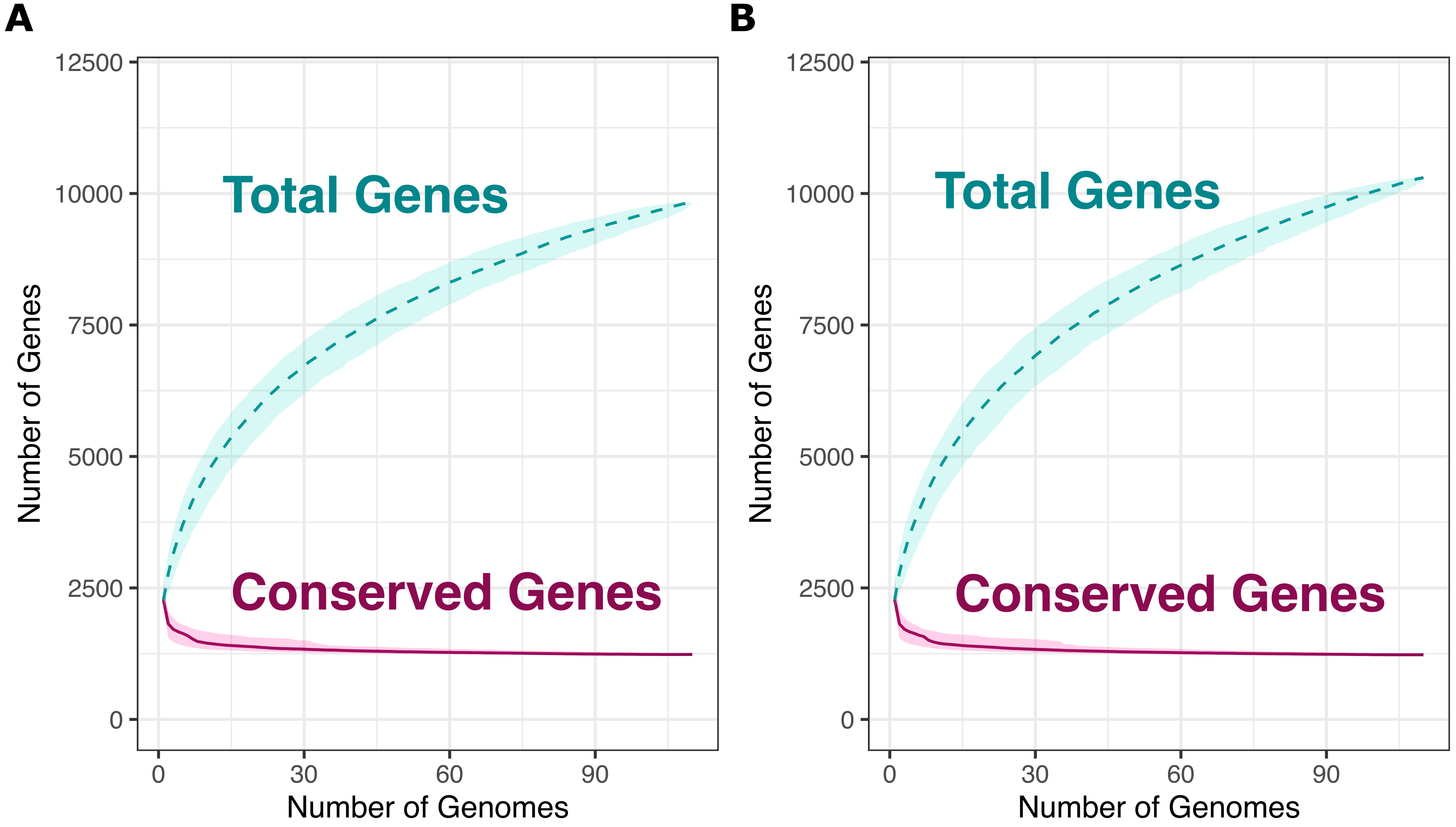


Supplementary Figure S3. Rarefaction curves for the Macrococcus pan- and core-genome, constructed using all 104 high-quality, publicly available Macrococcus genomes, plus six bovine-associated South African genomes sequenced here (n = 110 total Macrococcus genomes). Curves were constructed using PEPPAN, using a core genome threshold of 95% and an amino acid identity threshold of (A) 20% and (B) 40%. Curves showcase the accumulation of pan genes ("Total Genes") and core genes ("Conserved Genes") using 1,000 random permutations. Dashed and solid curved lines denote median values for pan and core genes, respectively, and shading surrounding each line denotes the respective 95% confidence interval.
