## Supplementary Figure S4 for "Genus-Wide Genomic Characterization of *Macrococcus*: Insights into Evolution, Population Structure, and Functional Potential"

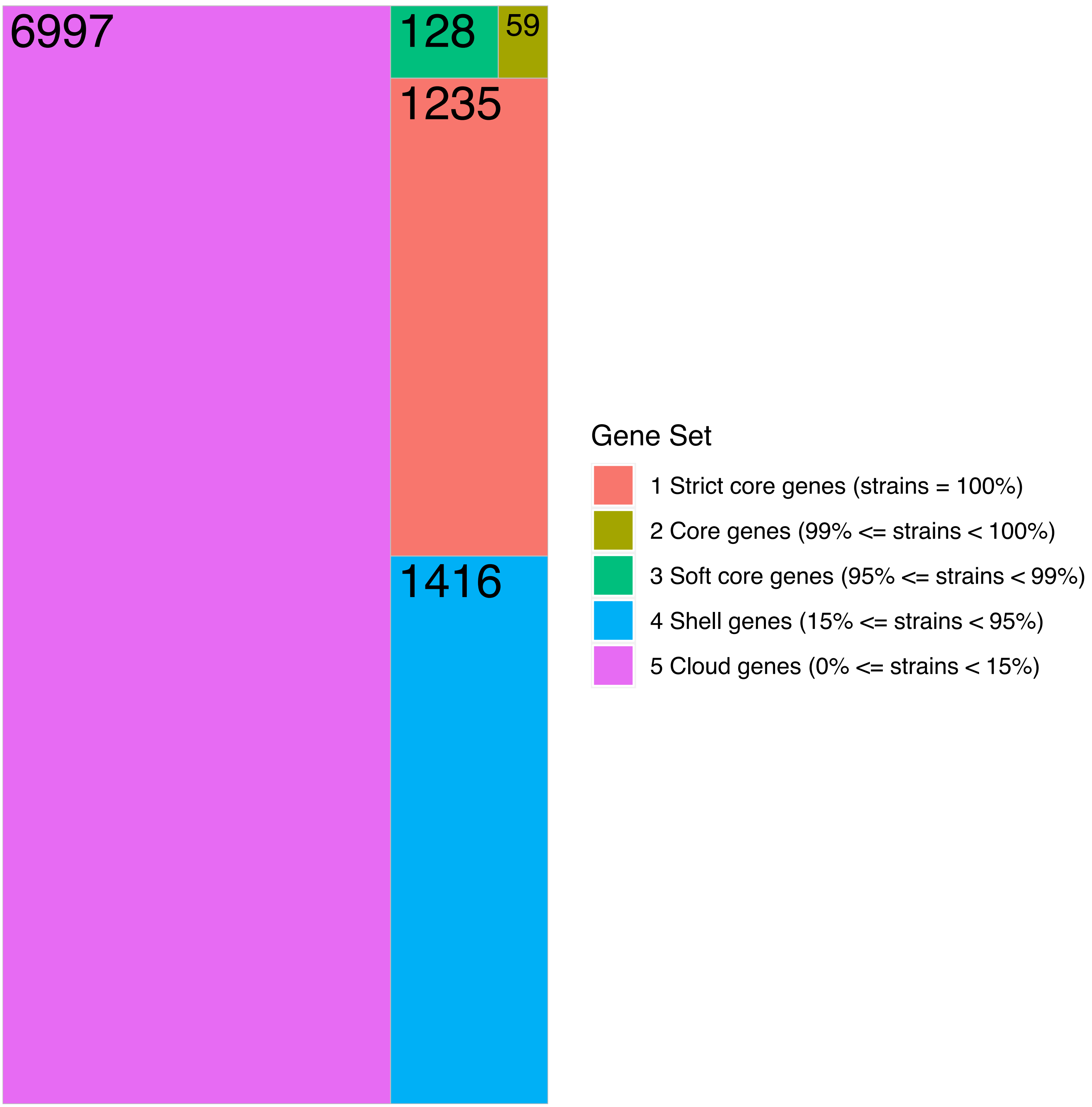


Supplementary Figure S4. Treemap showcasing the number of genes detected within a given percentage of Macrococcus genomes (out of 110 total genomes). Tile sizes are proportional to the number of genes detected within a given percentage of Macrococcus genomes; numerical labels within each tile denote the corresponding number of genes. PEPPAN was used to construct the core- and pan-genomes using a 20% amino acid identity threshold and a core genome threshold of 95%
