## Supplementary Figure S7 for "Genus-Wide Genomic Characterization of *Macrococcus*: Insights into Evolution, Population Structure, and Functional Potential"

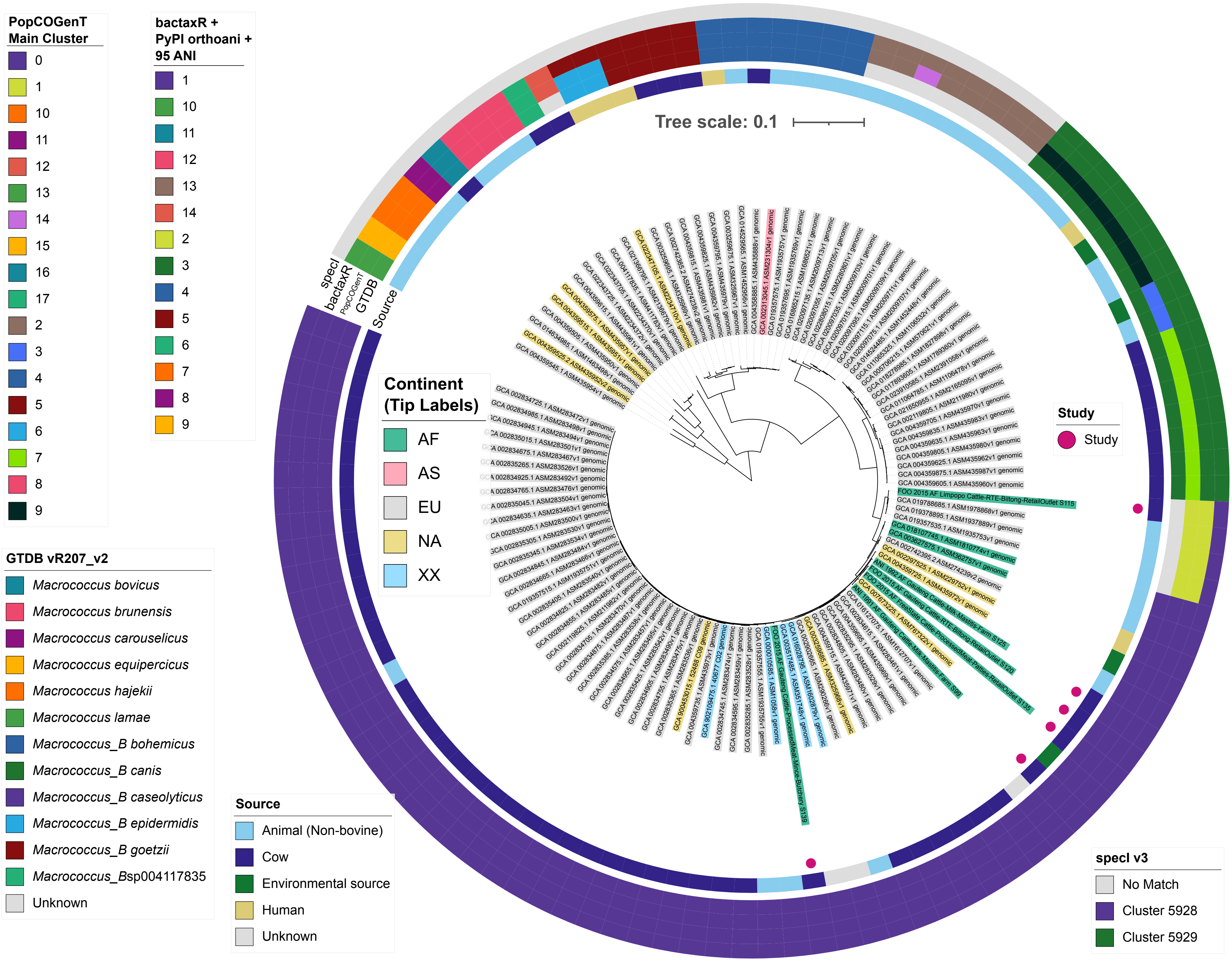


Supplementary Figure S7. Maximum likelihood (ML) phylogeny constructed using an amino acid multiple sequence alignment (MSA) of 120 bacterial marker genes detected in 110 Macrococcus genomes, plus the genome of Staphylococcus aureus str. DSM 20231 as an outgroup. The Genome Taxonomy Database Toolkit (GTDB-Tk) was used to construct the MSA. The tree was rooted using the outgroup (omitted for readability), and branch lengths are reported in substitutions per site. Tip label colors denote the continent from which each strain was reportedly isolated ("Continent"). Pink circles denote genomes sequenced in this study ("Study"). Rings surrounding the phylogeny denote (from interior to exterior) (i) the isolation source reported for each strain ("Source"), as well as species assignments obtained using four different taxonomic frameworks: (ii) Genome Taxonomy Database (GTDB) species, assigned using the Genome Taxonomy Database Toolkit (GTDB-Tk) v2.1.0 and GTDB vR207_v2 ("GTDB"); (iii) PopCOGenT "main clusters" (i.e., gene flow units, which attempt to mirror the classical species definition for animals and plants; "PopCOGenT"); (iv) genomospecies clusters delineated de novo using average nucleotide identity (ANI) values calculated via OrthoANI, bactaxR, and a 95 ANI genomospecies threshold (i.e., the threshold largely adopted by the microbiological community; "bactaxR"); (v) marker gene-based species clusters within the specI v3 taxonomy ("specI"). AF, Africa; AS, Asia; EU, Europe; NA, North America; XX, unknown/unreported geographic location.
