## Supplementary Figure S8 for "Genus-Wide Genomic Characterization of *Macrococcus*: Insights into Evolution, Population Structure, and Functional Potential"

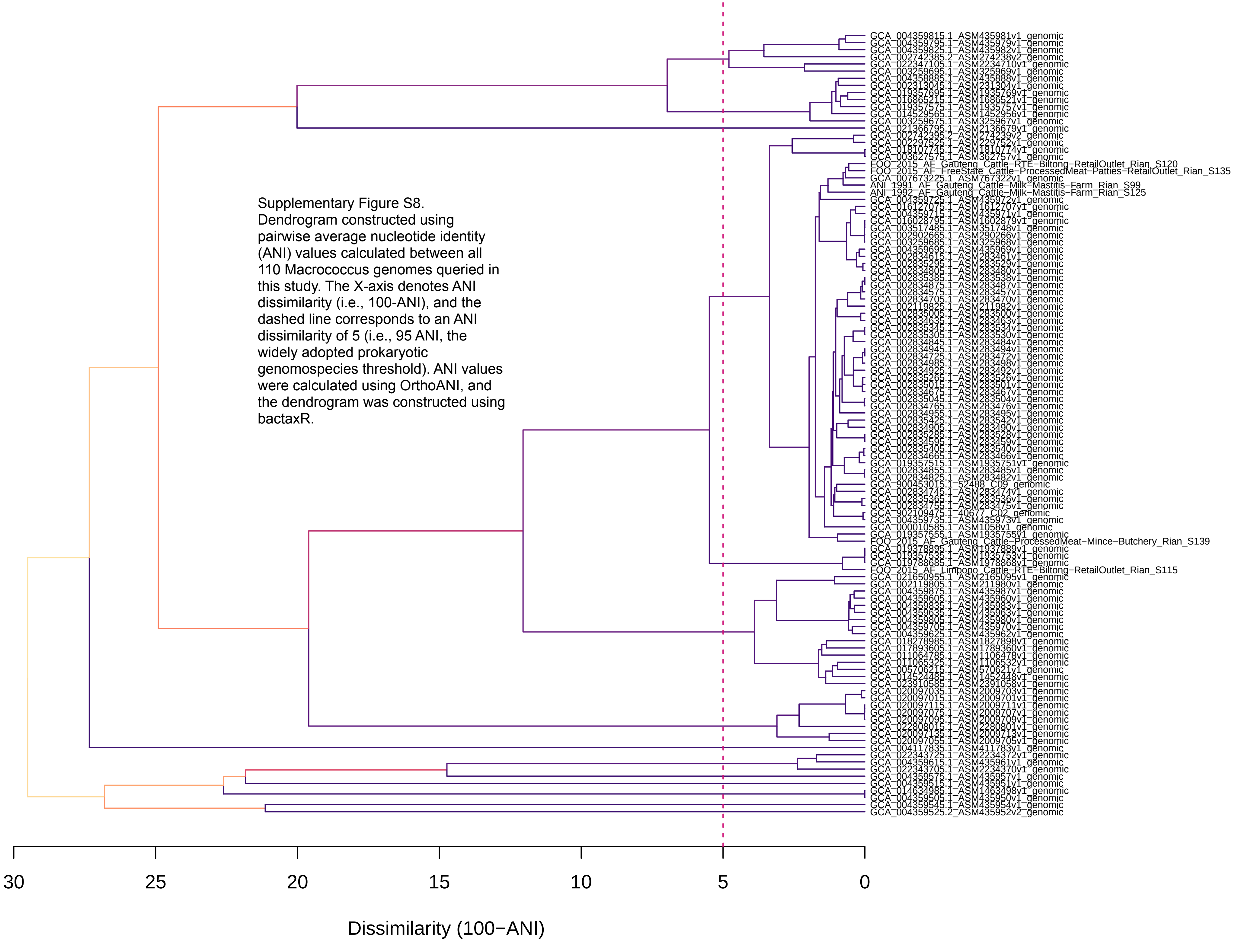


Supplementary Figure S8. Dendrogram constructed using pairwise average nucleotide identity (ANI) values calculated between all 110 Macrococcus genomes queried in this study. The X-axis denotes ANI dissimilarity (i.e., 100-ANI), and the dashed line corresponds to an ANI dissimilarity of 5 (i.e., 95 ANI, the widely adopted prokaryotic genomospecies threshold). ANI values were calculated using OrthoANI, and the dendrogram was constructed using bactaxR.
