## Supplementary Figure S9 for "Genus-Wide Genomic Characterization of *Macrococcus*: Insights into Evolution, Population Structure, and Functional Potential"

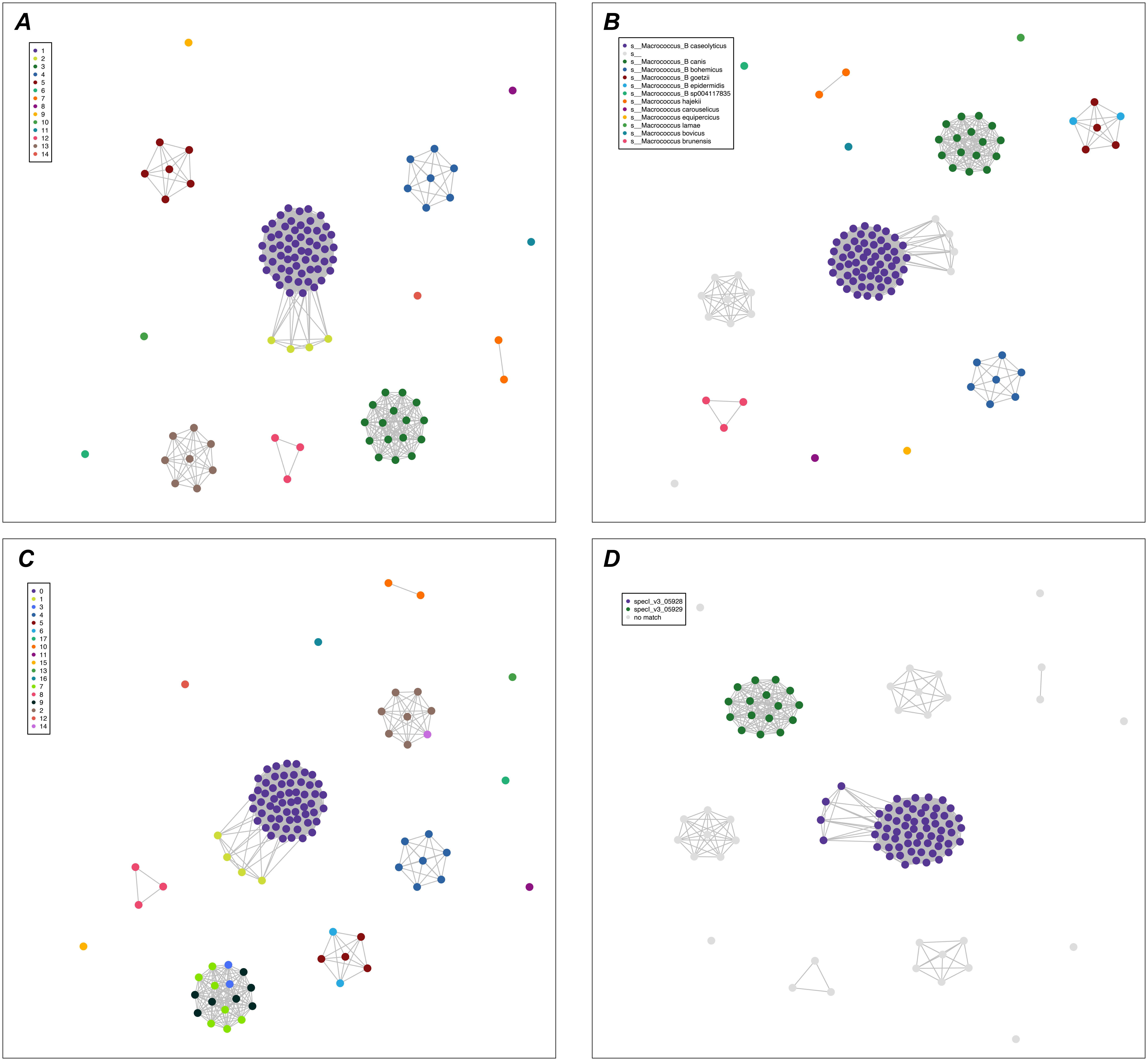


Supplementary Figure S9. Network constructed using pairwise average nucleotide identity (ANI) values calculated between 110 Macrococcus genomes. Nodes represent individual genomes, colored by the following: (A) genomospecies clusters delineated de novo using ANI values calculated via OrthoANI, bactaxR, and a 95 ANI genomospecies threshold (i.e., the threshold largely adopted by the microbiological community); (B) Genome Taxonomy Database (GTDB) species, assigned using the Genome Taxonomy Database Toolkit (GTDB-Tk) v2.1.0 and GTDB vR207_v2; (C) PopCOGenT "main clusters" (i.e., gene flow units, which attempt to mirror the classical species definition for animals and plants); (D) marker gene-based species clusters within the specI v3 taxonomy. Two nodes (genomes) are connected if they share at least 95 ANI with each other (calculated using OrthoANI). Networks were constructed and displayed using the ANI.graph function in bactaxR (default settings).
