## Supplementary Figure S10 for "Genus-Wide Genomic Characterization of *Macrococcus*: Insights into Evolution, Population Structure, and Functional Potential"

Supplementary Figure S10. Maximum likelihood (ML) phylogeny of all 104 high-quality, publicly available Macrococcus genomes, plus six bovine-associated South African genomes sequenced here (n = 110 total Macrococcus genomes). Tip label colors correspond to genomospecies assignments obtained via the Genome Taxonomy Database Toolkit (GTDB-Tk) v2.1.0 and GTDB vR207_v2. Genomes of strains isolated and sequenced in this study are denoted by pink circles (“Study”). Heatmaps to the right of the phylogeny denote (from left to right): (i) the source from which each strain was reportedly isolated (“Source”); (ii) the continent from which each strain was reportedly isolated (“Continent”); (iii) percentage of virulence factors (VF) present in the Virulence Factor Database (VFDB) core database, which were detected in each genome using DIAMOND blastp, with minimum amino acid identity and subject coverage thresholds of 40 and 50%, respectively (“VFDB VF”). The ML phylogeny was constructed using an alignment of 649 core genes identified among all 110 Macrococcus genomes, plus the genome of Staphylococcus aureus str. DSM 20231 (outgroup genome; NCBI RefSeq Assembly accession GCF_001027105.1), using Panaroo and a 50% protein family sequence identity threshold. The tree was rooted using the outgroup (omitted for readability), and branch lengths are reported in substitutions per site. Branch labels correspond to branch support percentages obtained using one thousand replicates of the ultrafast bootstrap approximation. AF, Africa; AS, Asia; EU, Europe; NA, North America; XX, unknown/unreported geographic location.

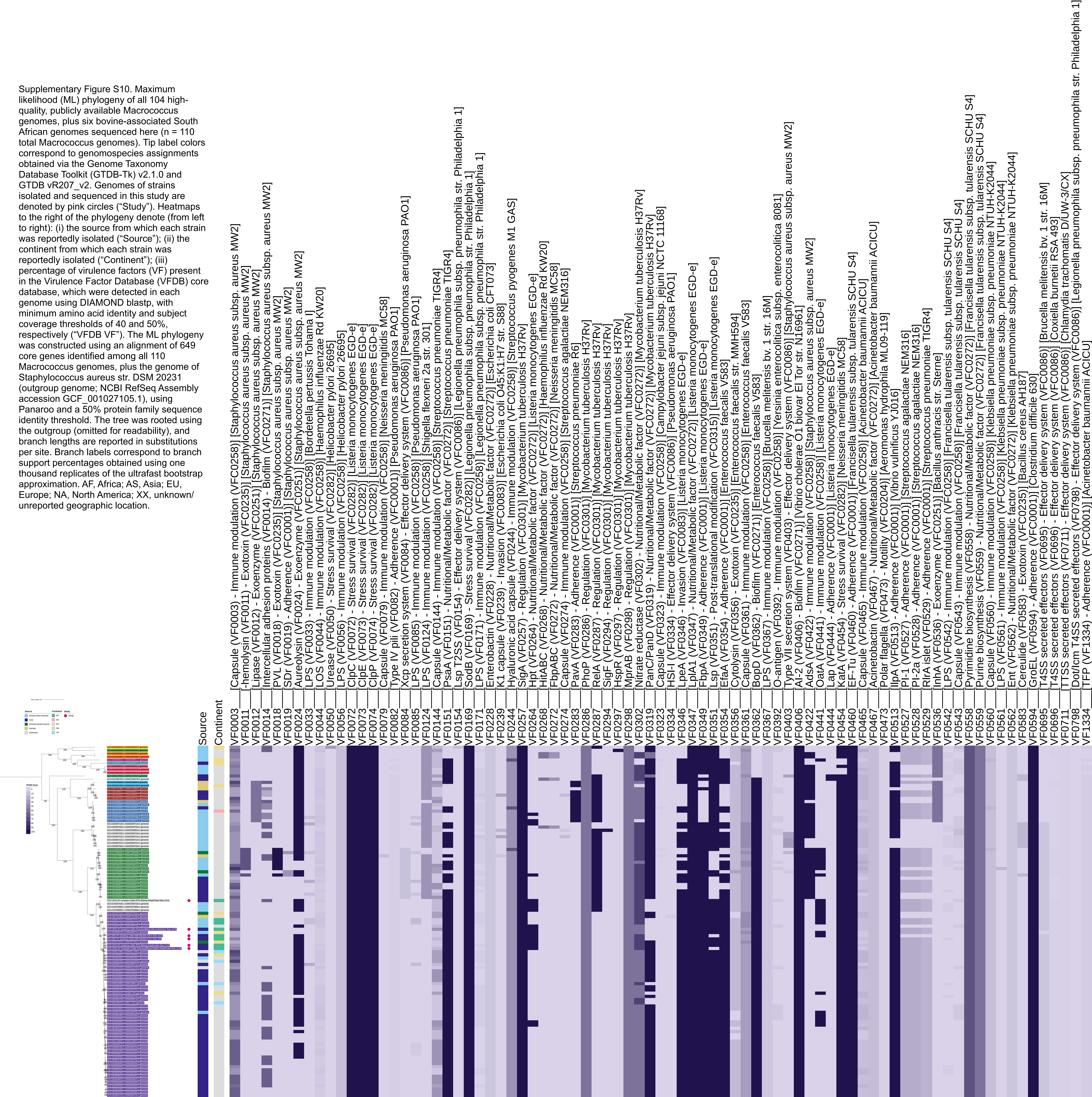
