## Supplementary Figure S11 for "Genus-Wide Genomic Characterization of *Macrococcus*: Insights into Evolution, Population Structure, and Functional Potential"

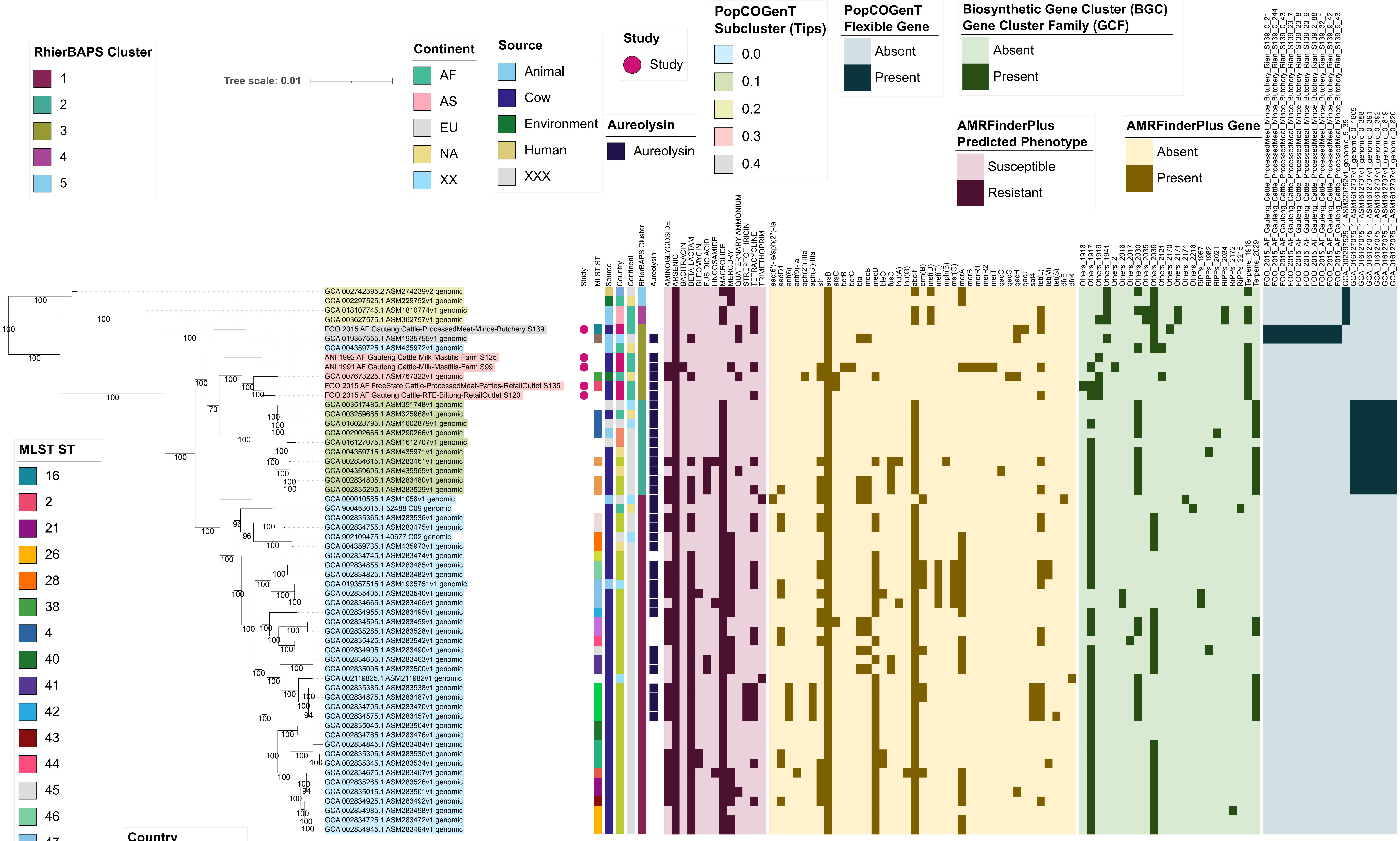


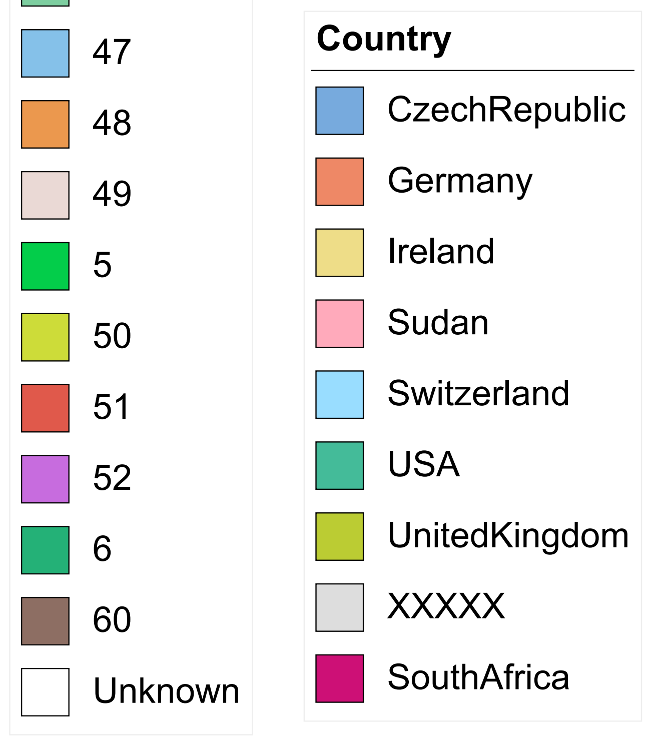


Supplementary Figure S11. Maximum likelihood (ML) phylogeny of 58 genomes assigned to the Genome Taxonomy Database's (GTDB) Macrococcus caseolyticus genomospecies. Tip label colors correspond to subcluster assignments obtained using PopCOGenT ("PopCOGenT Subcluster"). Pink circles denote genomes sequenced in this study ("Study"). Color strips/heatmaps to the right of the phylogeny denote (from left to right): (i) the sequence type (ST) assigned to each strain using in silico multi-locus sequence typing (MLST; "MLST ST"); (ii) the source from which each strain was reportedly isolated ("Source"); (iii) the country from which each strain was reportedly isolated ("Country"); (iv) the continent from which each strain was reportedly isolated ("Continent"); (v) the cluster assigned using RhierBAPS ("RhierBAPS Cluster"); (vi) presence of gene(s) sharing homology to aureolysin at 40% amino acid identity and 50% coverage ("Aureolysin"); (vii) predicted antimicrobial resistance (AMR) and stress response phenotype, obtained using AMR and stress response determinants identified via AMRFinderPlus ("AMRFinderPlus Predicted AMR Phenotype"); (viii) AMR and stress response determinants identified in each genome using AMRFinderPlus ("AMRFinderPlus Gene"); (ix) biosynthetic gene cluster (BGC) gene cluster families (GCFs) present in each genome ("Biosynthetic Gene Cluster [BGC] Gene Cluster Family [GCF]"); (x) presence and absence of flexible genes identified via PopCOGenT ("PopCOGenT Flexible Gene"). The ML phylogeny was constructed using an alignment of 1,751 core genes identified among all 58 Macrococcus caseolyticus genomes, plus an outgroup Macrococcus spp. genome from bactaxR Cluster 2 (NCBI GenBank Assembly accession GCA_019357535.1; Figure 1), using Panaroo and a 70% protein family sequence identity threshold. The tree was rooted using the outgroup (omitted for readability), and branch lengths are reported in substitutions per site. Branch labels correspond to branch support percentages obtained using one thousand replicates of the ultrafast bootstrap approximation. AF, Africa; AS, Asia; EU, Europe; NA, North America; XX, unknown/unreported geographic location; XXX, unknown/unreported source; XXXXX, unknown/unreported country.
