## Supplementary Figure S12 for "Genus-Wide Genomic Characterization of *Macrococcus*: Insights into Evolution, Population Structure, and Functional Potential"

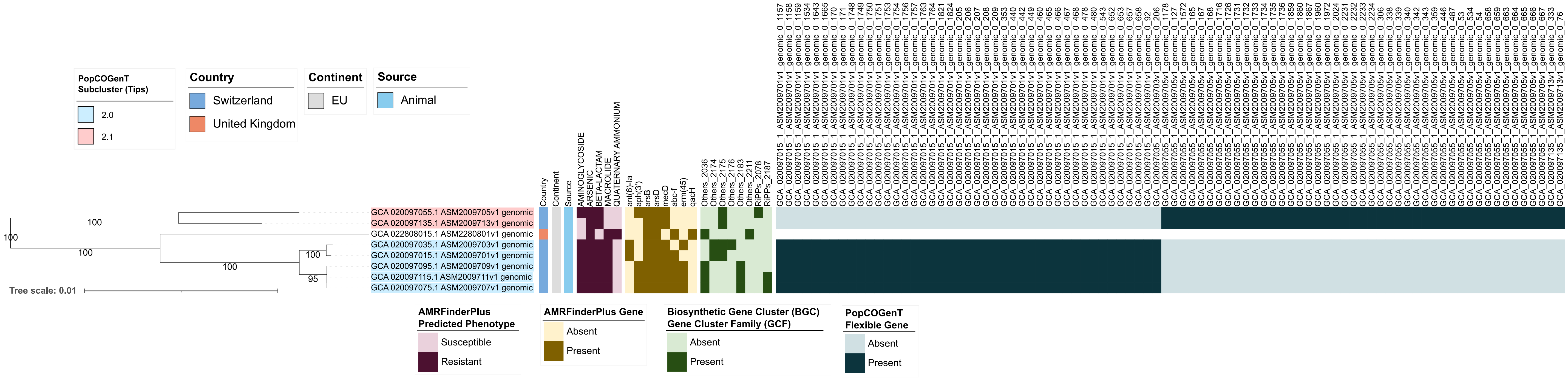


Supplementary Figure S12. Maximum likelihood (ML) phylogeny of eight genomes assigned to bactaxR Cluster 13 (i.e., Macrococcus armenti, based on average nucleotide identity [ANI]-based comparisons to species type strain genomes; Figure 1). Tip label colors correspond to subcluster assignments obtained using PopCOGenT ("PopCOGenT Subcluster"; one genome was not assigned to the same main cluster via PopCOGenT, and thus is not colored). Color strips/heatmaps to the right of the phylogeny denote (from left to right): (i) the country from which each strain was reportedly isolated ("Country"); (ii) the continent from which each strain was reportedly isolated ("Continent"); (iii) the source from which each strain was reportedly isolated ("Source"); (iv) predicted antimicrobial resistance (AMR) and stress response phenotype, obtained using AMR and stress response determinants identified via AMRFinderPlus ("AMRFinderPlus Predicted AMR Phenotype"); (v) presence and absence of AMR and stress response determinants identified in each genome using AMRFinderPlus ("AMRFinderPlus Gene"); (vi) biosynthetic gene cluster (BGC) gene cluster families (GCFs) identified in each genome ("Biosynthetic Gene Cluster [BGC] Gene Cluster Family [GCF]"); (vii) presence and absence of flexible genes identified via PopCOGenT ("PopCOGenT Flexible Gene"; for gene descriptions, see Supplementary Table S9). The ML phylogeny was constructed using an alignment of 1,416 core genes identified among all eight Macrococcus armenti genomes, plus an outgroup Macrococcus canis genome (NCBI GenBank Assembly accession GCA_014524485.1; Figure 1), using Panaroo and a 70% protein family sequence identity threshold. The tree was rooted using the outgroup (omitted for readability), and branch lengths are reported in substitutions per site. Branch labels correspond to branch support percentages obtained using one thousand replicates of the ultrafast bootstrap approximation. EU, Europe.
